# β-Nicotinamide mononucleotide: a novel broad-spectrum CRISPR inhibitor

**DOI:** 10.1101/2025.11.22.685598

**Authors:** Tong Wei, Wei Shen, Wenjun Li, Yingjie Song, Yunzhi Fa, Jing An, Yansong Sun, Hao Li

## Abstract

CRISPR–Cas systems have revolutionized genome editing with their precision and versatility, enabling transformative applications in various fields, especially in the treatment of genetic diseases. However, the clinical translation of this technology is hindered by challenges such as off-target effects and uncontrolled nuclease activity. At the same time, it has the possibility of causing biosecurity risks, underscoring the urgent need for reliable regulatory tools. Existing CRISPR inhibitors, primarily anti-CRISPR protein or exogenously synthesized small molecules, are limited by their specificity or bioavailability and long research period, unable to address the diverse CRISPR nucleases used in research and therapy. Based on the phenomena obtained from various in vitro and cell experiments, combining molecular dynamics simulation and bio - layer interferometry (BLI) analysis, here we report a naturally occurring small-molecule β-nicotinamide mononucleotide (NMN), the first known endogenous metabolite with broad-spectrum inhibitory activity against multiple CRISPR-associated proteins (Cas9, Cas12, and Cas13) through various mechanisms. Our findings establish NMN as a dual-purpose tool, which reduces cell damage caused by gene editing and mitigates risks of unintended genetic modifications in research and clinical settings. This discovery further shortens the distance between basic medicine and translational medicine, providing a new approach for developing endogenous regulatory molecules in genome engineering.

## Introduction

The CRISPR–Cas system, a bacterial adaptive immune mechanism repurposed for genome editing [1], has revolutionized biotechnology since its seminal demonstration in 2012 [2]. This RNA-guided nuclease technology enables precise modification of DNA and RNA across diverse species. Its transformative potential spans gene therapy for monogenic disorders, such as sickle cell anemia [3], as well as other broad areas [4∼8]. However, realization of its clinical promise is hindered by two critical limitations, namely, uncontrolled post-editing nuclease activity and off-target modifications. Persistent Cas protein activity can induce prolonged DNA damage, chromosomal rearrangements, and unintended edits in nontarget tissues, as observed in preclinical models where residual Cas9 causes insertional mutagenesis in diverse kinds of cells [9,10,11]. Recently approved CRISPR therapies, such as the ex vivo edited stem cell treatment Lyfgenia, also lack intrinsic mechanisms to terminate nuclease activity after editing, which may suffer from off-target effects due to CRISPR-related genetic information has been integrated into the host genome by a lentiviral vector. This critical gap creates an urgent demand for a clinically compatible inhibitor capable of broadly regulating diverse Cas nucleases while leveraging endogenous safety profiles [12]. In addition, vector-based CRISPR delivery systems, including lentiviruses and adenoviruses, carry risks of malicious use to harm key gene sites in the organism, which in extreme scenarios could lead to large-scale genetic perturbations in organisms. These challenges underscore the urgent need for robust blocker tools to modulate CRISPR activity with high precision [13].

Current CRISPR inhibitory strategies, including phage-derived anti-CRISPR (Acr) proteins [14,15,16] and synthetic small molecules [17], suffer from their specificity and organism repulsiveness. For example, AcrIIC1 and AcrIIA3 are two kinds of anti-Cas proteins and they target only Cas9 protein[18]; AcrVA4 targets only L. bacterium (Lb) and M. bovoculi (Mb) Cas12a but not Ac. species (As) Cas12a[19]; and AcrVIA1 targets only Cas13a[20], they mainly binding to the functional domains of the target protein, thereby inhibiting its biological activity, howevere as exogenous biomacromolecules, they perhaps produce an immune response in the body. Synthetic small-molecule inhibitors may exhibit poor cellular permeability or cytotoxicity, long R & D cycle, and possible body repulsiveness, which impede clinical translation. Due to their limited functionality and the difficulty of application, such inhibitors unable to address the expanding CRISPR toolbox, which now includes Cas12 and Cas13 for some new purposes.

Nicotinamide mononucleotide (NMN) is an endogenous metabolite derived from vitamin B3. It serves as a key precursor in the biosynthesis of nicotinamide adenine dinucleotide (NAD+), a coenzyme essential for cellular energy metabolism and genome maintenance. The therapeutic potential of NMN is amplified by its favorable pharmacokinetic profile. As an endogenous molecule, it exhibits high bioavailability when administered orally, with clinical studies confirming safety and tolerability in humans [21]. In previous studies, the connection between NMN and gene editing has mainly focused on the gene repair–promoting effect of NMN after its conversion into NAD+ as a precursor [22–26], here, we reported for the first time that NMN can bind to nucleic acids and Cas proteins directly, which confers inhibitory activity on multiple CRISPR-mediated gene editing. Further mechanistic investigations were conducted, revealing two CRISPR inhibition pathways. This means that we can harness its endogenous properties to develop safer, more effective strategies for maintaining genome integrity across gene therapy based on CRISPR genome-editing technology.

## Results

### 1. Inhibition of cell death after gene damage

Since NAD+ promotes the repair of genes, in order to verify whether there are changes in NAD+ levels within cells during the CRISPR-Cas9 mediated gene editing process, we continuously monitored the NAD+ levels of HEK-293 cells after knocking out the EMX1 gene by plasmid. The results indicated that, compared to normal cells, the NAD+ levels in the experimental group of cells decreased by 30.25 ±5.27%, 47.56±9.69%, and 62.98±2.03% at 12, 24, and 48 hours respectively after the gene knockout (Fig. 1A). To rule out the correlation of this phenomenon with EMX1 itself, we attempted to construct a CRISPR–Cas9–induced gene damage model that involves widespread damage to the gene by targeting the highly repeatable Alu sequence widely present on a variety of animal chromosomes (Fig. 1B). The results showed that compared to normally cultured cells, after Alu was knocked out, the cells still showed 69.82 ± 3.89% decreasing NAD+ levels after 48 hours (Fig. 1C), these data show that the decrease in NAD+ levels may be a common manifestation of all CRISPR-Cas9 system-mediated gene-edited cells.

**Fig. 1.**
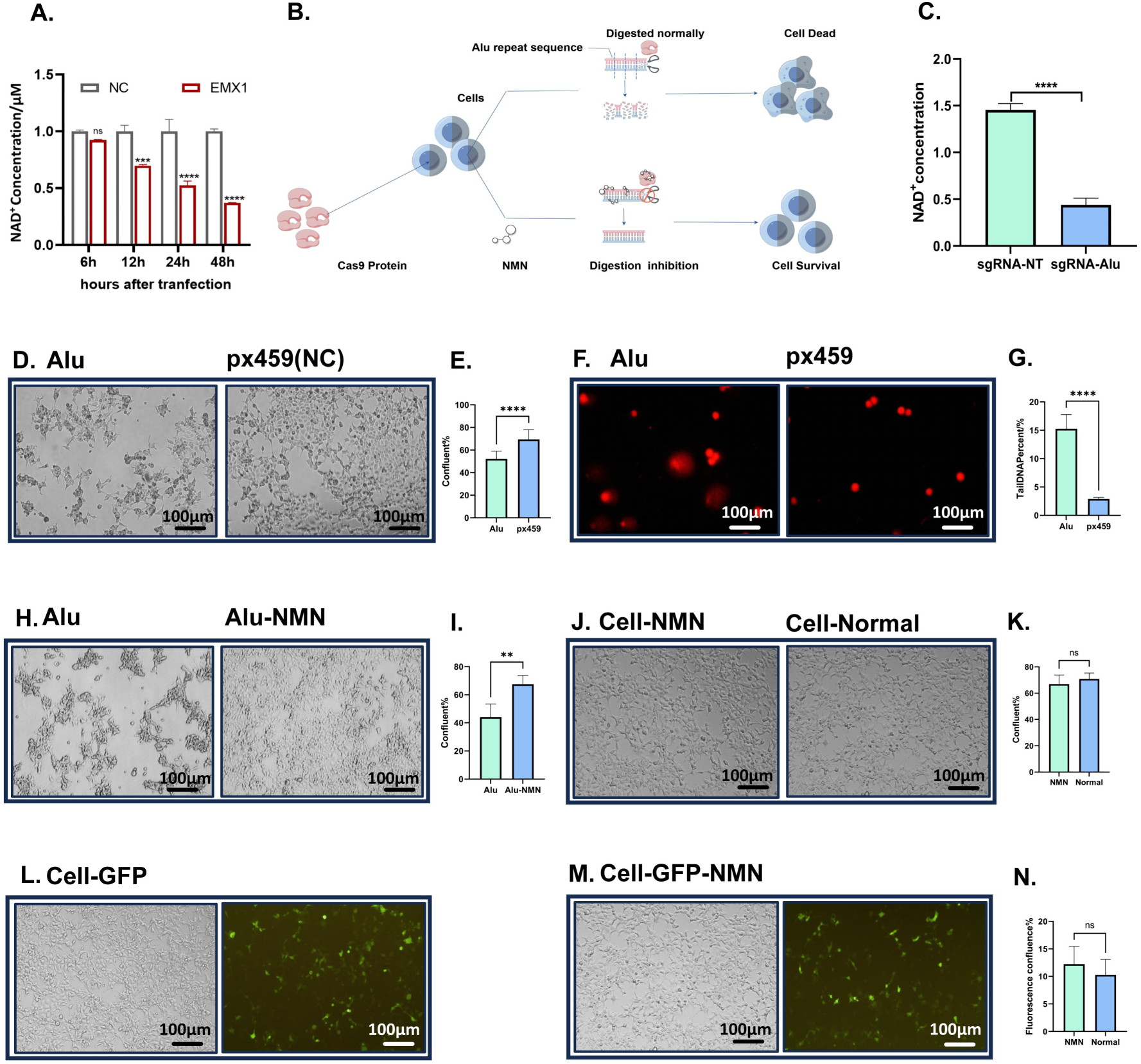
Protective effect of NMN on CRISPR-induced cell death due to gene damage. A. Changes in cellular NAD+ levels after single-target knockout. B. Principle of CRISPR–Cas9 targeting highly repetitive sequences, leading to cellular gene damage. C. Changes in cellular NAD+ levels after the knockout of Alu repetitive sequences for 48 hours. D. Comparison of cell confluence with and without transfection of CRISPR–Cas9 system plasmids targeting the Alu sequence. E. Statistical data of cell confluence with and without transfection of CRISPR–Cas9 system plasmids targeting the Alu sequence. F. Comet experiments with and without transfection of CRISPR–Cas9 system plasmids targeting the Alu sequence. G. Statistical data on the ratio of comet tail length to total comet length in the comet assay. H. Comparison of cell confluence with and without NMN addition after transfection of CRISPR–Cas9 system plasmids targeting the Alu sequence. I. Statistical data of cell confluence with and without NMN addition after transfection of CRISPR–Cas9 system plasmids targeting the Alu sequence. J. Comparison of cell confluence with and without NMN addition. K. Statistical data of cell growth at 12 hours before and after NMN addition. L. Normal transfection of cells. M. Transfection of cells with the addition of NMN at the time of the transfection. N. Statistical data of the transfection efficiency before and after the addition of NMN.

As for the construction of gene damage model, compared with the control group in which nontargeting plasmids were transfected into HEK-293 cells, the cell confluence of the experimental group transfected with the CRISPR–Cas9 plasmid targeting Alu decreased by 17.18 ± 3.215% (Fig. 1D, E). The tailing of the genome after electrophoresis revealed that the average ratio of the comet tail length of the cell genome to the total comet size in the Alu-targeting group was 12.35±2.277% higher than that in the control group (Fig. 1F, G), these indicate that following the transfection of a CRISPR–Cas9 system targeting Alu sequences into the cells, gene damage occurs, which is accompanied by a certain degree of cell death.

Subsequently, since NMN is an important supplement and precursor to NAD+, to verify whether NMN supplementation effect on cell death mediated by CRISPR–Cas9 system,we supplemented NMN into the culture medium while delivering the CRISPR–Cas9 system, observing the confluence of cells before and after delivery; then excluding NMN’s potential impact on cell growth rate and transfection efficiency. When NMN was added during transfection, the cell confluence increased by 23.57±5.088% after 48 hours (Fig. 1H, I). We further showed that there was no significant change in cell growth rate (Fig. 1J, K) and transfection efficiency (Fig. 1L, M, N) after the addition of NMN. These data indicate that the addition of NMN can reduce cell death mediated by the CRISPR–Cas9 system and that this effect should not be attributed to the interference with transfection efficiency or an increase in cell growth rate. This finding suggests that NMN may inhibit the damage caused by the CRISPR–Cas9 system to cellular genes.

### 2. Inhibition of in vitro digestion caused by CRISPR

To eliminate confounding variables in cellular experiments and confirm whether NMN directly interacts with the CRISPR–Cas9 system, we further employed the CRISPR–Cas9 in vitro cleavage system for subsequent validation. NMN was coincubated or separately preincubated with Cas9 protein and substrate nucleic acid before the restriction enzyme system starts working, substrates were then subjected to electrophoretic analysis to observe the changes in cleavage efficiency. The Cas9 reaction system normally cleaves the target nucleic acids into two distinct fragments. However, only the uncut substrate band was observed following electrophoresis after NMN coincubation with the whole Cas9 system (Fig. 2A) and preincubation with substrate or Cas9 proteins (Fig. 2B, C), and the cut bands as in the positive control group, which perform the enzyme digestion reaction with standard system, appeared in the acidic control group (Fig. 2A, B, C). Specifically, in exploring NMN coincubation with the whole Cas9 system, Therefore, the results showed that the enzymatic cleavage efficiency of Cas9 was weakened regardless of whether NMN was coincubated with the Cas9 reaction system or preincubated with substrates or proteins, and this phenomenon was not related to acidity. Notably, unlike the distinct band pattern observed when NMN was coincubated with the complete system, the preincubation with substrate condition resulted in a smeared band pattern accompanied by substantial retention within the sample loading well, while the extent of nucleic acids cleavage inhibition resulting from protein preincubation, as well as that observed when NMN was coincubated with the entire reaction mixture, only led to a small amount of nucleic acid depositing at the sample wells.

**Fig. 2.**
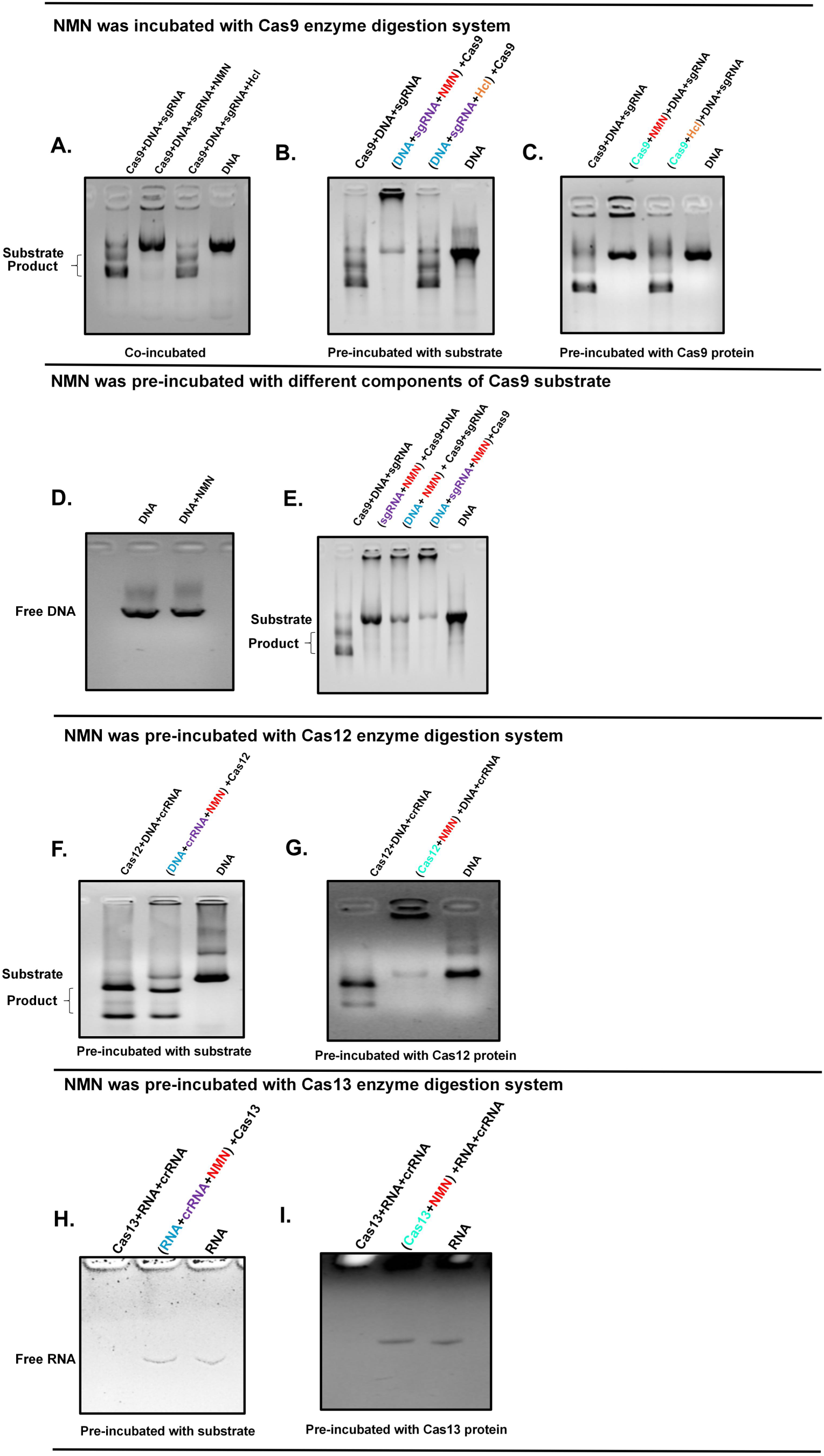
Inhibition of CRISPR enzyme digestion efficiency by NMN and the preincubation of NMN and substrate or Cas9. A. The cleavage efficiency of the CRISPR–Cas9 in vitro digestion kit was inhibited after adding NMN to the system, while the acidic control group was digested normally. B. The cleavage efficiency of the CRISPR–Cas9 in vitro digestion kit was inhibited after NMN was preincubated with substrate nucleic acids, and there was different deposition of nucleic acids at the wells. C. The cleavage efficiency of the CRISPR–Cas9 in vitro digestion kit was inhibited after NMN was preincubated with Cas9, and there was different deposition of nucleic acids at the wells. D. Direct incubation of DNA with NMN did not cause electrophoretic retardation. E. The cleavage efficiency of the CRISPR–Cas9 in vitro digestion kit was inhibited after NMN was preincubated with sgRNA, target DNA or both of them in substrate nucleic acids, and there was different deposition of nucleic acids at the wells. F. The cleavage efficiency of the CRISPR–Cas12 in vitro digestion kit was inhibited after NMN was preincubated with the corresponding substrate nucleic acids. G. The cleavage efficiency of the CRISPR–Cas12 in vitro digestion kit was inhibited after NMN was preincubated with Cas12. H. The cleavage efficiency of the CRISPR–Cas13 in vitro digestion kit was inhibited after NMN was preincubated with the corresponding substrate nucleic acids. I. The cleavage efficiency of the CRISPR–Cas13 in vitro digestion kit was inhibited after NMN was preincubated with Cas13.

To further investigate the reasons for this occurrence, we added NMN separately to sgRNA, target DNA or both of them in substrate, then perform pre-incubation before enzyme digestion and electrophoresis. Compared with the electrophoresis bands of the single substrate, incubation of the substrate solely with NMN did not result in changes to the shape, size, intensity or position (Fig. 2D). However, when sgRNA, target DNA or both of them in substrate were separately preincubated with NMN before enzyme digestion, the electrophoresis bands of substrate after enzymatic digestion showed some changes in intensity (Fig. 2E). These results indicate that NMN induces changes in intensity and nucleic acid depositing electrophoretic banding patterns after incubation with sgRNA, target DNA or both of them in substrate. In summary, result above suggesting that NMN interacts differently with sgRNA, target DNA or both of them in substrate and Cas9 protein within the Cas9 reaction system. When NMN is coincubated with substrates and proteins, NMN may preferentially interact with proteins and inhibit cleavage efficiency.

To explore whether this inhibition phenomenon is applicable to other common CRISPR digestion systems, we added NMN to Cas12 and Cas13 systems for incubation and digestion. The Cas12 reaction system cleaves the target nucleic acids into two distinct fragments similar to the Cas9 reaction system. However, only the uncut substrate band was observed following electrophoresis after NMN preincubation with substrate or Cas12 proteins (Fig. 2F, G). The Cas13 system completely cleaves RNA, resulting in the absence of any substrate bands in the electrophoresis lane. However, when NMN was preincubated with its substrate or Cas13 proteins, the bands of the substrate appeared normally. The above results indicate that the addition of NMN to the Cas12 and Cas13 cleavage systems also demonstrates varying levels of inhibition in cutting efficiency. Therefore, this substance can act on multiple CRISPR systems, showing the broad-spectrum nature of its inhibitory effects.

### 3. Inhibition efficiency in distinct conditions

To further delineate the conditions under which NMN inhibits CRISPR systems, we employed the CRISPR–Cas9 system as a representative model. We systematically titrated the concentration of NMN (20mM, 40mM, 80mM) and varied its preincubation period (10min, 25min, 35min) with the substrate complex or Cas9 portein. Cleavage inhibition efficiency was then accurately quantified using quantitative PCR (qPCR) by designed qPCR primers on both sides of the cleavage ends of the substrate DNA (Fig. 3A). The results showed that when NMN concentration was varied during a fixed 30-minute preincubation with the substrate complex (Fig. 3B, C, D), the cleavage efficiency of Cas9 in positive control reactions averaged 93.27%; 80 mM NMN reduced Cas9 cleavage efficiency by a mean of 42.5%; 40 mM NMN reduced cleavage efficiency by a mean of 29.5%; and 20 mM NMN reduced cleavage efficiency by a mean of 10.1%. When preincubation time was varied at a fixed NMN concentration of 40 Mm (Fig. 3E, F, G), positive control cleavage efficiency averaged 94.3%; 35-minute NMN incubation reduced Cas9 cleavage efficiency by a mean of 40.0%; 25-minute incubation reduced cleavage efficiency by a mean of 20.2%; and 10-minute incubation reduced cleavage efficiency by a mean of 2.9%. This indicates that the CRISPR inhibition caused by NMN preincubation with substrate is incubation time–dependent and concentration-dependent.We also found that preincubation of NMN with the Cas9 protein rapidly achieved a mean CRISPR–Cas9 system inhibition efficiency of 68.7% (Fig. 3H, I, J, K, L, M), even at reduced concentrations and with minimal incubation time. This indicates that NMN and protein preincubation have stronger inhibition efficiency against the CRISPR system, which is consistent with the results of previous electrophoresis.

**Fig. 3.**
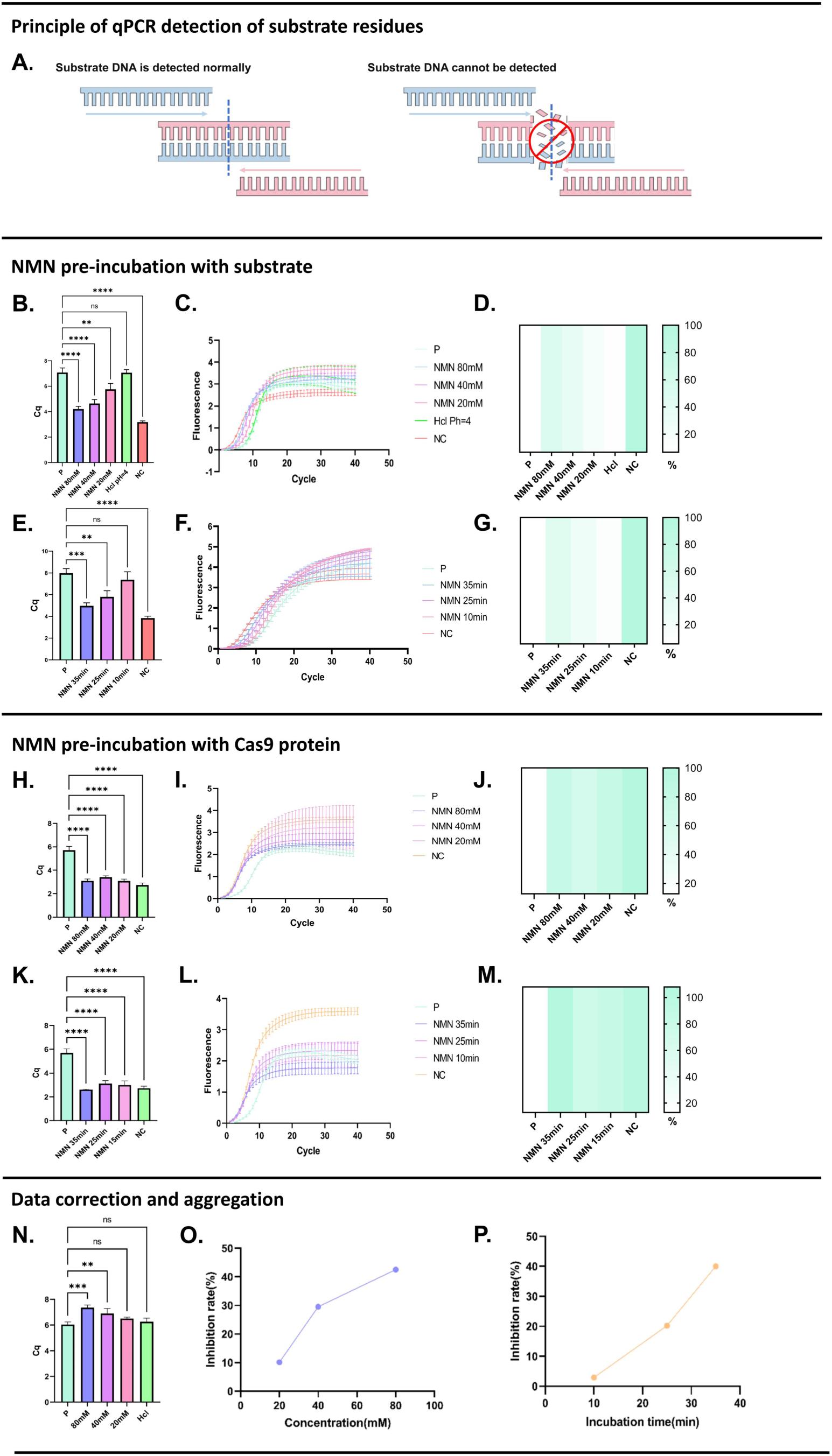
Accurate evaluation of the inhibition efficiency of the CRISPR–Cas9 system by the preincubation of NMN and substrate. A. Principle of qPCR detection of substrate residues. B. Cq value statistics chart of the results of qPCR detection of residual substrate after 30-minute preincubation of different concentrations of NMN with the corresponding substrate of the CRISPR–Cas9 system. C. The results of qPCR detection of residual substrate after 30-minute preincubation of different concentrations of NMN with the corresponding substrate of the CRISPR–Cas9 system. D. Heat map of the results of qPCR detection of residual substrate percentage after 30-minute preincubation of different concentrations of NMN with the corresponding substrate of the CRISPR–Cas9 system. E. Cq value statistics chart of the results of qPCR detection of residual substrate after different preincubation times of 40 mM NMN with the corresponding substrate of the CRISPR–Cas9 system. F. The results of qPCR detection of residual substrate after different preincubation times of 40 mM NMN with the corresponding substrate of the CRISPR–Cas9 system. G. Heat map of the results of qPCR detection of residual substrate percentage after different preincubation times of 40 mM of NMN with the corresponding substrate of the CRISPR– Cas9 system. H. Cq value statistics chart of the results of qPCR detection of residual substrate after 30-minute preincubation of different concentrations of NMN with Cas9. I. The results of qPCR detection of residual substrate after 30-minute preincubation of different concentrations of NMN with Cas9. J. Heat map of the results of qPCR detection of residual substrate percentage after 30-minute preincubation of different concentrations of NMN with Cas9. K. Cq value statistics chart of the results of qPCR detection of residual substrate after different preincubation times of 40 mM of NMN with Cas9. L. The results of qPCR detection of residual substrate after different preincubation times of 40 mM NMN with Cas9. M. Heat map of the results of qPCR detection of residual substrate percentage after different preincubation times of 40 mM of NMN with Cas9. N. The different detection efficiency of qPCR after adding different concentrations of NMN to the Product to be tested of qPCR system with the same amount of substrate. O. The relationship between the inhibition efficiency of CRISPR–Cas9 in vitro digestion and the concentration of NMN after 30-minute preincubation with NMN and corresponding substrate of the CRISPR–Cas9 system. P. The relationship between the inhibition efficiency of CRISPR–Cas9 in vitro digestion and the preincubation time of 40 mM NMN and corresponding substrate of the CRISPR–Cas9 system.

To detect the impact of NMN on the qPCR detection system, we added different concentrations of NMN to the uncut substrate, consistent with the dosage used during preincubation, and we observed the detection results. The qPCR analysis revealed a reduction rate of 59.96%, 45.05%, 28.06%, in concentration of added NMN is 80mM, 40mM, 20mM in assay detection efficiency upon NMN supplementation (Fig. 3N). Consequently, this NMN-associated effect requires consideration in the interpretation of subsequent inhibition efficiency data. Accounting for this, the true magnitude of cleavage inhibition is likely greater than the measured values.

### 4. Exploration of the preliminary mechanism

To investigate the underlying mechanism, we focused on the CRISPR–Cas9 system. Electrophoretic Mobility Shift Assay (EMSA) was employed to assess the binding capacity of the NMN-preincubated substrate complex to the Cas9 protein and the binding capacity of the NMN-preincubated Cas9 protein to the substrate complex. The results indicated that when the substrate was preincubated with NMN before incubation with the dCas9 protein, the bands of the substrate DNA exhibited a significant lag compared with free DNA, the same mobility lag distance as when the substrate was normally incubated with the dCas9 protein (Fig. 4A). In contrast, when the dCas9 protein was preincubated with NMN before being incubated with the substrate, the corresponding substrate bands did not exhibit a lag phenomenon (Fig. 4B). This demonstrates that after the preincubation of NMN with the substrate, the dCas9 protein continues to bind to the substrate normally, but the NMN-preincubated dCas9 protein fails to effectively bind the substrate. This binding impairment strongly suggests divergent inhibitory mechanisms arising from NMN interactions with either the protein or the substrate. Specifically, NMN preincubated with the substrate may primarily suppresses CRISPR activity by inhibiting the cleaving activity function of the Cas protein. Conversely, NMN preincubated with the Cas protein predominantly disrupts activity by impairing its molecular recognition and substrate-binding capability.

**Fig. 4.**
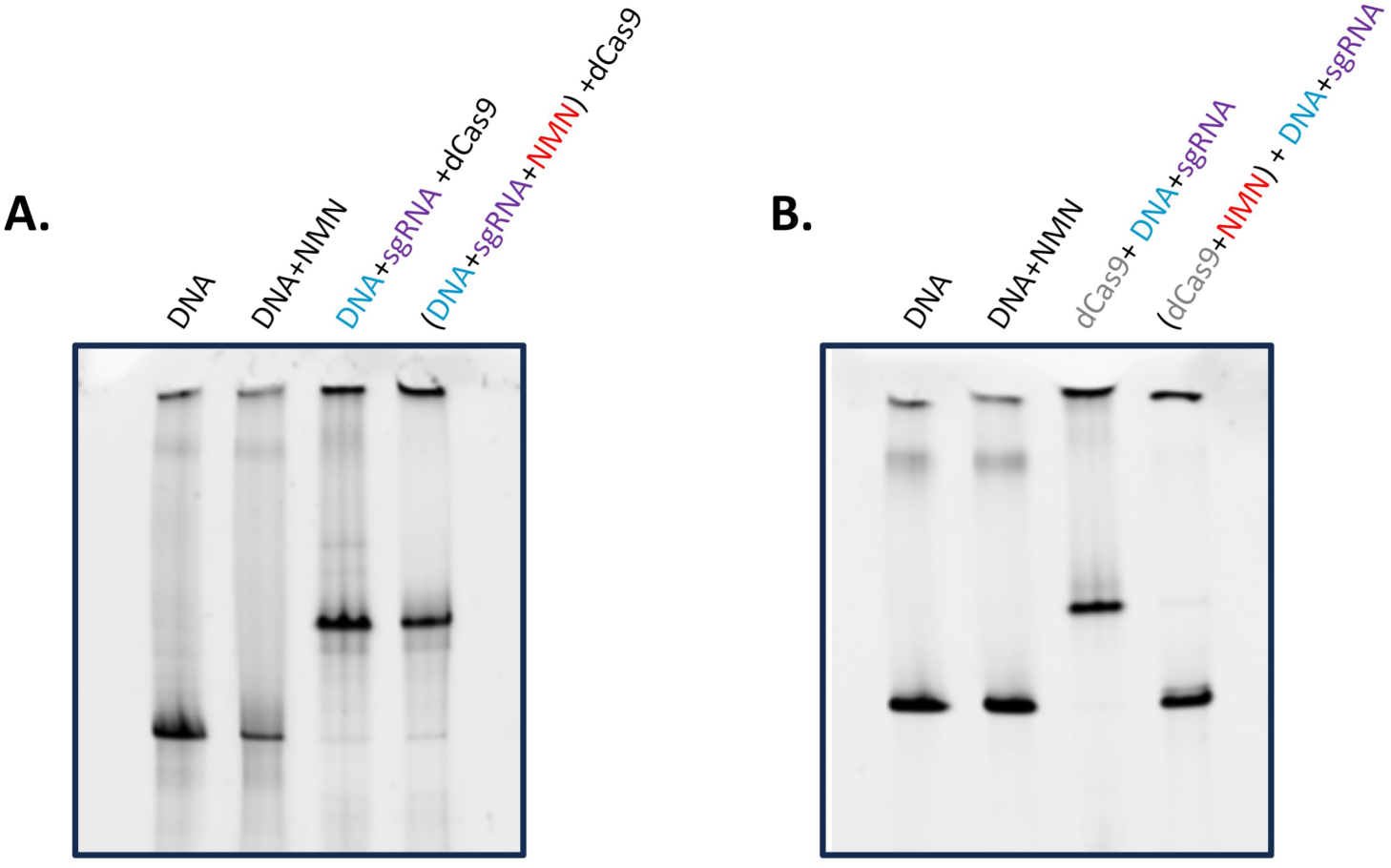
EMSA test result of the inhibition principle of the CRISPR–Cas9 system by the preincubation of NMN and substrate or by the preincubation of NMN and Cas9. A. EMSA test result of the preincubation of NMN and substrate. B. EMSA test result of the preincubation of NMN and Cas9.

### 5. Different mechanisms of action

Previous results suggest that NMN may have a connection with the Cas9 protein or its substrate. To verify this hypothesis, we employed Bio-Layer Interferometry (BLI analyses) to validate the differential binding affinity of compound NMN toward the two components. The result revealed that the binding signal of nucleic acids or Cas9 protein was enhanced with the enhancement of the concentration of added ligand NMN (Fig. 5A, B), this suggests that NMN exhibits concentration-dependent binding to both nucleic acids and Cas9 protein. Crucially, the dissociation constant KD for NMN binding to Cas9 was 14.7-times greater than that for NMN–nucleic acid interaction (Fig. 5C, D). The KD ratio reflects the ligand concentration required to occupy half-maximal binding sites, thereby demonstrating that NMN binds Cas9 with substantially higher affinity than nucleic acids. These results explain why the electrophoretic characteristics of NMN coincubated with the Cas9 enzyme cutting system are closer to those of NMN preincubated with the Cas9 protein. Namely, compared with the substrate, NMN preferentially binds to the protein.

**Fig. 5.**
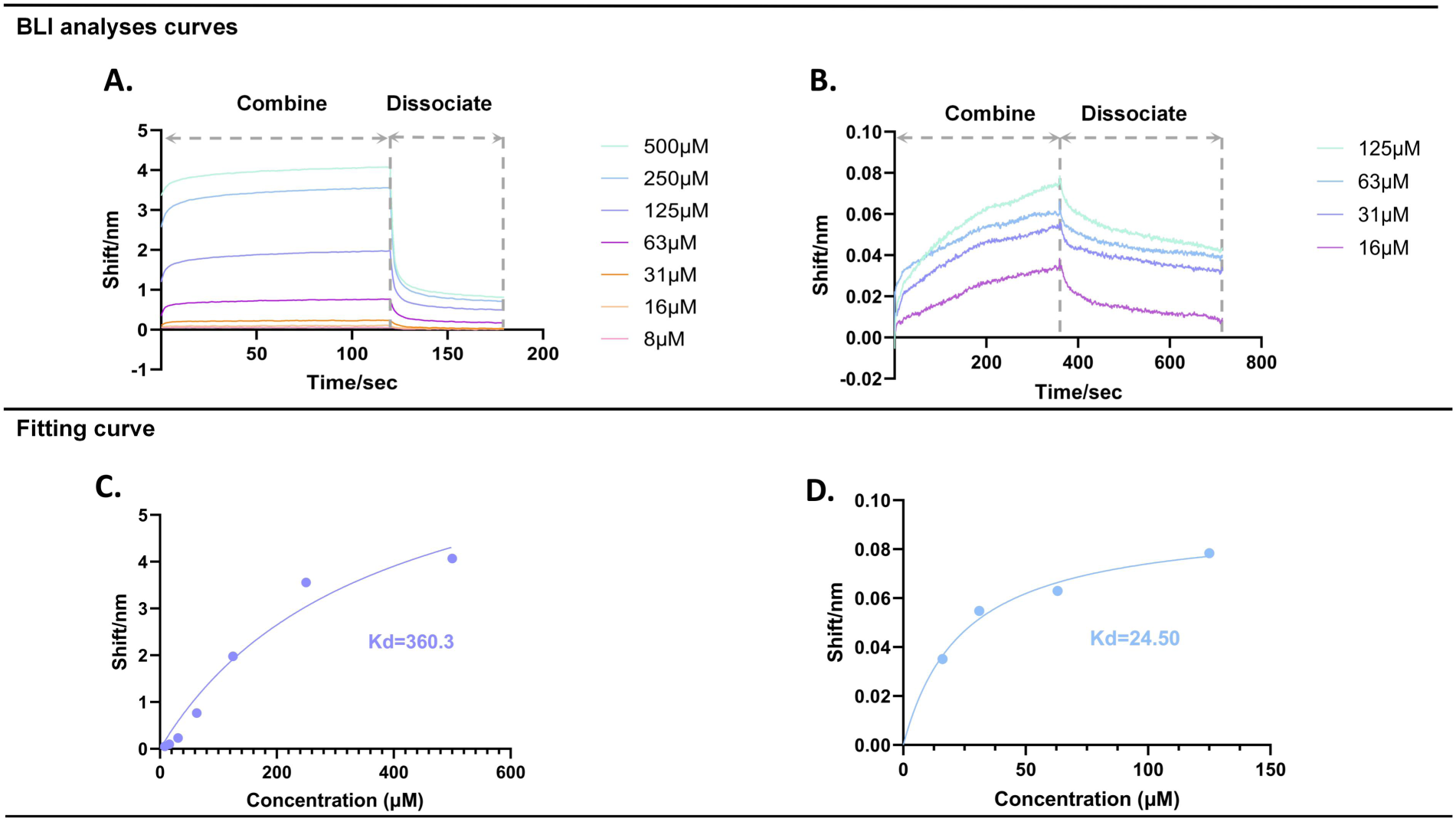
BLI analyses data of NMN with the Cas9 protein and DNA. A. BLI analyses curves of different concentrations of NMN and DNA. B. BLI analyses curves of different concentrations of NMN and Cas9 protein. C. Affinity analysis data fitting curves of NMN and DNA. D. Affinity analysis data fitting curves of NMN and Cas9 protein.

Subsequently, to explore the specific mode of action of NMN with nucleic acids or proteins, we employed molecular dynamics (MD) simulations to investigate the interactions between NMN and the substrate DNA, the NMN-preincubated substrate and the Cas9 protein, and NMN and the Cas9 protein. The first simulations revealed that NMN forms 342 hydrogen bonds with 263bp dsDNA, which includes 57.6% nonstandard hydrogen bonds, resulting in binding. Furthermore, the calculated binding free energy indicated a weak association between NMN and the substrate (Fig. 6A–G).

**Fig. 6.**
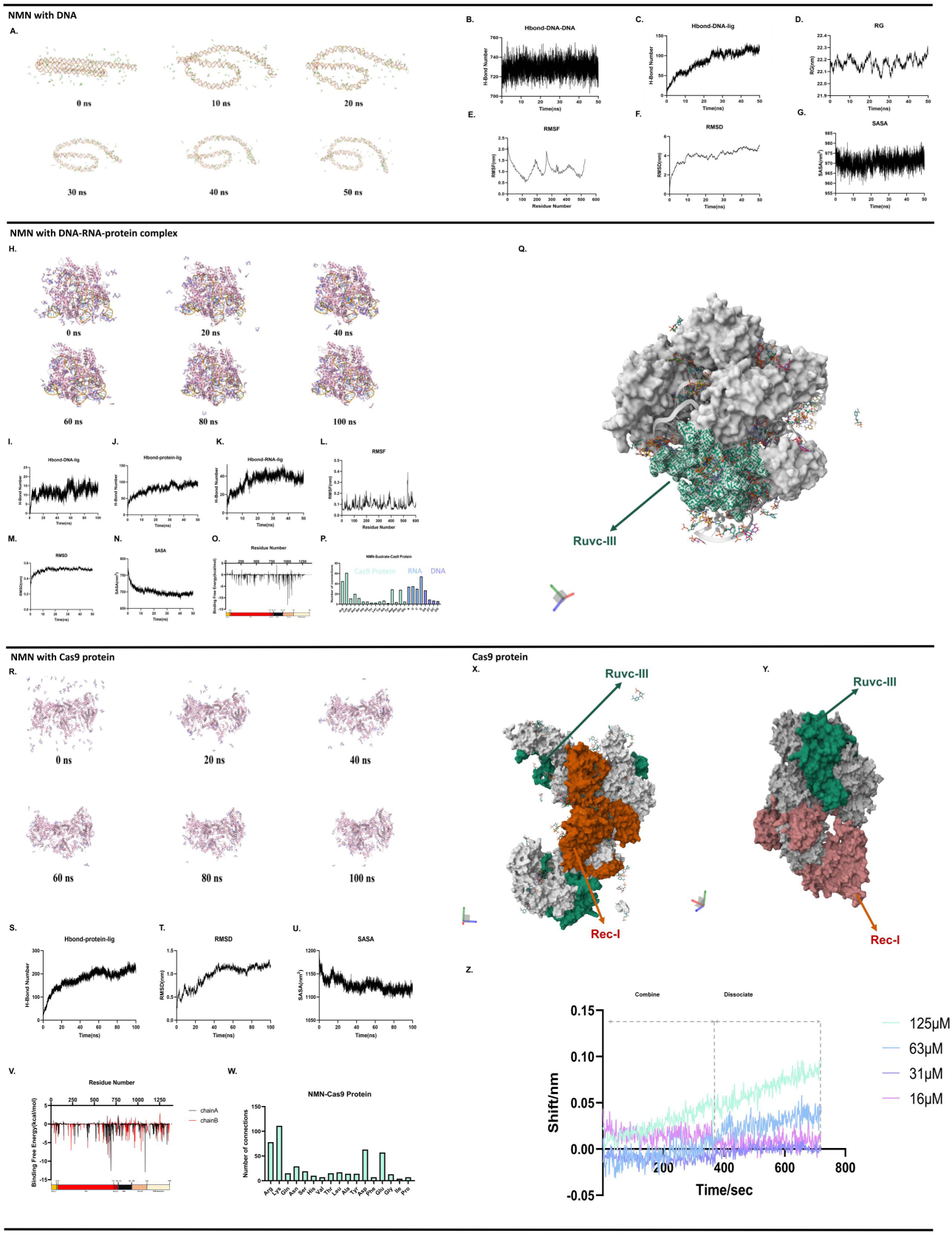
Molecular dynamics (MD) simulation results of nucleic acids, DNA–RNA–protein complex, and Cas9 protein with NMN. A. Snapshot of nucleic acids incubation with NMN. B. Number of DNA–DNA hydrogen bonds in the MD result of nucleic acids incubation with NMN. C. Number of DNA–NMN hydrogen bonds in the MD result of nucleic acids incubation with NMN. D. Radius of gyration of DNA in the MD process of nucleic acids incubation with NMN. E. Root mean square fluctuations of DNA in the MD process of nucleic acids incubation with NMN. F. Root mean square deviation of DNA in the MD process of nucleic acids incubation with NMN. G. Solvent accessible surface area of DNA in the MD process of nucleic acids incubation with NMN. H. Snapshot of DNA–RNA–protein complex incubation with NMN. I. Number of DNA–NMN hydrogen bonds in the MD result of DNA–RNA–protein complex incubation with NMN. J. Number of Cas9–NMN hydrogen bonds in the MD result of DNA–RNA–protein complex incubation with NMN. K. Number of sgRNA–NMN hydrogen bonds in the MD result of DNA–RNA–protein complex incubation with NMN. L. Root mean square fluctuations of DNA in the MD process of DNA–RNA–protein complex incubation with NMN. M. Root mean square deviation of DNA in the MD process of DNA–RNA–protein complex incubation with NMN. N. Solvent accessible surface area of DNA in the MD process of DNA–RNA–protein complex incubation with NMN. O. Statistics of Cas9 binding domain with NMN in the MD process of DNA–RNA–protein complex incubation with NMN. P. Statistics of binding sites of DNA–RNA–protein complex incubation with NMN. Q. Main binding site of DNA–RNA–protein complex incubation with NMN. R. Snapshot of the Cas9 protein incubation with NMN. S. Number of Cas9–NMN hydrogen bonds in the MD result of the Cas9 protein incubation with NMN. T. Root mean square deviation of the Cas9 protein in the MD process of the Cas9 protein incubation with NMN. U. Solvent accessible surface area of the Cas9 protein in the MD process of the Cas9 protein incubation with NMN. V. Statistics of Cas9 binding domain with NMN in the MD process of the Cas9 protein incubation with NMN. W. Statistics of binding sites of the Cas9 protein incubation with NMN. X. Main binding site of the Cas9 protein in two binding methods (Direction 1). Y: Main binding site of the Cas9 protein in two binding methods (Direction 2). Z.BLI analyses curves of NMN and knockout Cas9 protein.

Further, we positioned NMN within the interfacial region between the substrate and the Cas9 protein in the Cas9–substrate complex. The MD simulations revealed that after 10 ns of simulation, the relative interaction frequency of NMN with protein, RNA, and DNA a ratio of 10:4:1. Following 100 ns of simulation, this ratio shifted to 20:13:3 (Fig. 6H). Specifically, the calculated binding free energy of NMN to Cas9 exceeded that to the RNA component, which in turn was greater than that to the DNA component. Consequently, the majority of NMN molecules migrated towards the Cas9 protein (Fig. 6I–N).

Computational analysis indicated that the migrated NMN primarily occupied the catalytic pocket within the Cas9 RuvC nuclease domain (Fig. 6O, Q), mainly dependent on hydrogen bonding. During the internal migration process, NMN binds to 15 kinds of amino acids of the Cas9 protein. It primarily binds with arginine, lysine, glutamic acid, and aspartic acid, accounting for 66.5% of the total amino acid bindings, while their number makes up only 25.8% of the total amino acids. Arginine and lysine constitute 43.3% of the total amino acid bindings, while their number makes up only 10.7% (Fig. 6P). This occupancy suggests that NMN impairs Cas9 cleavage efficiency by sterically occluding the enzyme’s catalytic site after preincubation with the substrate.

After simulating the incubation of NMN alone with Cas9, we found that NMN can efficiently bind to the Rec I and Ruvc domains (Fig. 6V, X), mainly dependent on hydrogen bonding and strong electrostatic interactions (Fig. 6R–U). According to the calculation results, there was no significant change in the affinity tendency of NMN to amino acids in the case of direct incubation between NMN and the Cas9 protein compared with the internal migration process (Fig. 6W). This observation contrasts the previous one: a large amount of NMN binds to the Rec I domain compared with the incubation of NMN with DNA–RNA–protein complex (Fig. 6X,Y), and the Rec I domain is primarily responsible for binding with sgRNA; therefore, this circumstance may lead to the failure of RNP complex formation, which is consistent with our EMSA experiment results.

To validate these simulation results, we knocked out the RuvC and RecI domains of the Cas9 protein and expressed it. Subsequently, the BLI analysis was performed again with the knockout proteins. The results demonstrated that the knockout proteins no longer exhibited affinity for NMN. This confirms that the RuvC and RecI domains indeed play a crucial role in the binding process between Cas9 protein and NMN.

## Discussion

The continuous advancement of third-generation gene editing technologies has transformed therapeutic gene editing from a theoretical concept into a burgeoning medical field commanding a huge market [27]. Nevertheless, this technology remains fraught with substantial risks, predominantly stemming from the uncontrollable nature of in vivo gene editing. Multiple research groups have proposed various mitigation strategies. However, existing solutions have been limited by suboptimal inhibitory efficacy or pronounced immunogenicity in humans. Our research group identified a natural, human-derived, broad-spectrum CRISPR inhibitor that demonstrated robust suppression against three major CRISPR systems commonly deployed in gene editing applications through different mechanisms of action.

In this study, we demonstrated that NMN delivery effectively suppressed cell death induced by CRISPR–Cas9 gene editing and speculated that NMN inhibited the cutting efficiency of the CRISPR system. This phenomenon was validated through well-controlled biochemical assays designed to minimize confounding variables, and it was subsequently extended to CRISPR–Cas12 and CRISPR–Cas13 systems. Mechanistic investigations employing multiple analytical approaches revealed that NMN inhibits CRISPR systems via distinct molecular interactions with core components. Specifically, the results of cell experiments, electrophoresis experiments, and qPCR quantification showed that NMN could interact with sgRNA, target DNA or both of them in substrate and Cas protein in the CRISPR system to inhibit the enzyme cleavage effect mediated by the CRISPR system. The BLI analysis and electrophoresis results showed that NMN was more inclined to interact with proteins when it played a role in the CRISPR system. The MD simulations, BLI analysis and EMSA results revealed that preincubation of NMN with CRISPR–Cas9 substrates resulted in NMN binding to nucleic acids, followed by migration toward the RuvC nuclease domain of the Cas9 protein after NMN-preincubated substrate complex binding to Cas9 proteins, thereby inhibiting the cleavage ability of Cas proteins. Under physiological conditions, NMN may remain bound to nucleic acids while connected with the Cas9 protein and facilitate the formation of a quaternary complex (protein–dsDNA–sgRNA–NMN). This hypothesis aligns with the results of gel electrophoresis. That is, after NMN was preincubated with the substrate, although the protein could bind to the substrate, the cleavage efficiency was reduced. When NMN was preincubated with sgRNA, NMN predominantly formed complexes with sgRNA-bound Cas9, reducing cleavage efficiency. In this situation, only a small amount of DNA bound to sgRNA was carried to form complexes, and unbound DNA substrates exhibited pronounced electrophoretic separation (intense bands). After NMN preincubation with DNA, NMN-bound DNA formed ternary complexes with the Cas9 protein, resulting in attenuated band intensity due to reduced free DNA. Following NMN preincubation with sgRNA and DNA, synergistic effects minimized free DNA, yielding the faintest electrophoretic bands. The results of the MD calculations and EMSA experiments revealed that preincubation of NMN with the Cas9 protein directly blocked substrate binding via interactions with the Rec I domain, which is mainly responsible for binding to sgRNA and then targeting the target DNA, thereby efficiently inhibiting target DNA cleavage. Thus, in the absence of substrate binding to the protein, the electrophoresis bands retained integrity, which corresponded to the results of electrophoresis.

Since its inception, CRISPR technology has been accompanied by severe technical risks and ethical issues. As CRISPR technology continues to develop and expand in application, our control over biological genetic information has become increasingly proficient. However, this has led to growing concerns regarding the biosafety issues triggered by this technology. Currently, we have identified several CRISPR inhibitors, yet their clinical development has been hampered by concerns related to their inhibition spectrum and safety. NMN, a new type of broad-spectrum inhibitor with good biological adaptability and diverse mechanisms of action, may provide assistance in this regard. While our study established NMN as a validated CRISPR inhibitor and elucidated its underlying mechanisms, in vivo delivery efficiency and consequent gene editing suppression efficacy require further investigation in subsequent experiments.

## Materials

Biological Safety Cabinet (art. No. NU-433-400S) was purchased from Nuaire, Inc.USA.Carbon Dioxide Incubator (art. No. MCO-170AICDL-PC) was purchased from Puhexi Co., Ltd. Centrifuge (art. No. 5702JL453003) was purchased from Eppendorf Ltd. Fluorescence Inverted Microscope (art. No. CKX53) was purchased from Olympus Corporation. SCILANS Adjustable Mixer (art. No. VB218XN0021453) was purchased from SCILOGEX Corporation (U.S.). MiniL-12G Mini Centrifuge (art. No. MiNiL-12G-240514003) was purchased from Shanghai Maigao Scientific Instruments Co., Ltd.PHS-3E PH Meter (art. No. 600721N0022010214) was purchased from Shanghai INESA Scientific Instrument Co., Ltd. Mini Gel Tank (art. No. 1104241850), and PCR Instrument Applied Biosystems (art. No. A24812) were purchased from Thermo Fisher Scientific Inc.PowerPac Basic (art. No. 041BR319570) was purchased from Bio - Rad Laboratories (Shanghai) Co., Ltd. LightCycler 96 Instrument(art. No. 05815916001) was purchased from Roche Group (Germany).

SpCas9 Nuclease (art. No. 32101), and1 0× HOLMES Buffer for Cas9(art. No. 32041), and TOLO Cas9 control target and SgRNA(art. No. 32020), and Cas9 control Target dsDNA (art. No. 32020-TP1),and Cas9 Control sgRNA (art. No.32020-TP2), and LbCas12a Nuclease (art. No.32108-01),and TOLO Cas12a control target and crRNA (art. No.32021-01),and 10×HOLMES Buffer for Cas12a (art. No.32108-03),and Cas12a Control Target DNA and crRNA (art. No.32021-01),and LwaCas13a Nuclease (art. No.32117-03),and Cas13a Control Target RNA and crRNA (art. No.32024-01),and TOLO Cas13a control target and crRNA (art. No.32024-01)were purchased from Tolo Biotech Co., Ltd.

DMEM basic (1×) (art. No. C11995500BT),and TrypLE^TM^ Expree (1×) (art. No. 12605),and LTX and Plus™ Reagent (art. No. 15338100),and NativePAGE^TM^ 20× Running Buffer (art. No. BN2001),and NativePAGE^TM^ 3-12% Bis-Tris Gel (art. No. BN1003BOX)were purchased from Thermo Fisher Scientific Inc.PBS (1×)(art. No. PB180327)was purchased from Wuhan Pricella Biotechnology Co., Ltd. Nuclease-free Water (art. No. BL510B) was purchased from Lanjieker Technology Co., Ltd. 50×TAE buffer (art. No. B1110) was purchased from BeiJing Applygen Technologies Inc.HiPure Agarose(art. No. MF-103-01) was purchased from Guangzhou Magen Biotechnology Co.,Ltd. M5 Gelred Plus Nucleic acids Stain (10000X) (art. No. MF079-plus-01) was purchased from Mei5 Biotechnology Co., Ltd. 2K Plus Ⅱ DNA Marker (art. No. BM121) was purchased from TransGen Biotech; 6×DNA loading Buffer (Blue/Cyan) (art. No. 1109566) was purchased from Tiangen Biotech (Beijing)Co., Ltd.2× Super Pfx Master Mix (art. No. CW2965M) was purchased from Jiangsu Cowin Biotechnology Co., Ltd.

## Methods

### 1 Cell gene damage test (Cell Culture and Transfection)

HEK293 cells were plated at a density of 1 × 10⁶ cells per well in 24-well plates. Cells were cultured in 1 mL of Dulbecco’s Modified Eagle Medium (DMEM) supplemented with 10% fetal bovine serum (FBS), 2 mM L-glutamine, 1%penicillin and streptomycin. And the seeded plates were incubated for 24 hours at 37°C in a humidified atmosphere containing 5% CO₂. Then transfecting cells with plasmids, and the Use of plasmids in DNA Damage Condition is the CRISPR-Cas9-Alu target plasmid; in control Condition is the px459 plasmid (empty vector control), at first configure the two solutions of the transfection system separately: Solution A: 50 µL Opti-DMEM, 1 µg plasmid DNA (See figure legends for specific plasmids), and 10 µL PLUS™ Reagent were combined and incubated at room temperature for 10 minutes. Solution B: 50 µL Opti-DMEM and 10 µL Lipofectamine™ LTX Reagent were combined and incubated at room temperature for 10 minutes. subsequently, solutions A and B were combined, gently mixed, and incubated at room temperature for an additional 10 minutes to allow complex formation. The complete transfection mixture (100 µL/well) was added dropwise across five different locations within each well containing cells and culture medium. Cells were then returned to the 37°C, 5% CO₂ incubator. Cell confluence was assessed and recorded via phase-contrast microscopy 48 hours post-transfection. Representative images were captured.

### 2 The comet test to confirms the cause of the reduced confluence of the cells

The primary agarose layer was prepared by melting 1% normal-melting-point agarose, pouring it into a casting tray, and sequentially immersing slides before excision of the lateral gel segments (removing the outer 1/5 width). Processed slides were air-dried overnight at room temperature for subsequent use. For cell preparation, adherent cells were trypsinized, washed once with PBS, and pelleted by centrifugation; the resulting pellet was resuspended in PBS to a density of 1×10⁵ cells/mL. Separately, 0.75% low-melting-point agarose was melted and maintained at 37°C in a water bath to prevent solidification. Cell-agarose composites were generated by gently mixing 30 μL cell suspension with 50 μL liquefied agarose, pipetting the mixture onto the center of the pre-coated slides, and covering with coverslips to exclude air bubbles, followed by 15-min solidification at 4°C in darkness. For lysis, slides were decoverslipped and immersed in pre-chilled lysis buffer at 4°C for 2 hr under light-protected conditions. Post-lysis slides underwent three sequential 1-min washes in distilled water with manual agitation (mechanical shaking prohibited). Washed slides were aligned parallel to the electric field in an electrophoresis chamber, submerged in pre-chilled alkaline electrophoresis buffer (pH>13), and incubated for 20 min at 4°C in darkness for DNA unwinding, with the chamber surrounded by an ice bath. Electrophoresis was performed at 20 V (300 mA) for 25 min under light protection. Neutralization was achieved through three 1-min washes in Tris-HCl buffer (0.4 M, pH 7.5) with manual handling to prevent gel detachment, followed by air-drying at room temperature. Staining was performed by applying 50 μL ethidium bromide solution (500× diluted) onto gels, immediately covering with coverslips for uniform distribution, and incubating for 10 min in darkness. Imaging was conducted immediately post-staining using fluorescence microscopy at 200× magnification to capture comet morphologies.

### 3 Cell gene damage inhibition test

HEK293 cells were plated in 24-well plates as previously described and incubated for 24 hours at 37°C under a humidified atmosphere containing 5% CO₂. Upon removal from the incubator, 500 µL of culture medium was aspirated and discarded from each well. Fresh medium was prepared as follows: DNA Damage Group: 500 µL fresh culture medium (DMEM supplemented with 10% FBS, L-glutamine, penicillin, and streptomycin) was added per well; NMN Treatment Group: 500 µL fresh culture medium containing 20 mM NMN was added per well, resulting in a final concentration of 10 mM NMN upon addition. Cells were transfected immediately following medium replacement using the CRISPR-Cas9-Alu plasmid construct, according to the transfection protocol detailed above. And cell confluence was assessed and representative phase-contrast microscopy images were captured at 48 hours post-transfection.

### 4 Exploration of the effect of NMN on cell growth efficiency

HEK293 cells were plated in 24-well plates as previously described and incubated for 24 hours at 37°C under a humidified atmosphere containing 5% CO₂. Upon removal from the incubator, 500 µL of culture medium was aspirated and discarded from each well. Fresh medium was prepared as follows: Untreated Control Group: 500 µL of fresh culture medium (DMEM supplemented with 10% FBS, L-glutamine, penicillin, and streptomycin) was added per well. NMN Treatment Group: 500 µL fresh culture medium containing 20 mM NMN was added per well, resulting in a final concentration of 10 mM NMN upon addition. And cell confluence was assessed and representative phase-contrast microscopy images were captured 12 hours post-medium replacement.

### 5 Exploration of the effect of NMN on transfection efficiency

HEK293 cells were plated in 24-well plates as previously described and incubated for 24 hours at 37°C under a humidified atmosphere containing 5% CO₂. Upon removal from the incubator, 500 µL of culture medium was aspirated and discarded from each well. Fresh medium was prepared as follows: Untreated Control Group: 500 µL of fresh culture medium (DMEM supplemented with 10% FBS, L-glutamine, penicillin, and streptomycin) was added per well. NMN Treatment Group: 500 µL fresh culture medium containing 20 mM NMN was added per well, resulting in a final concentration of 10 mM NMN upon addition. Cells were transfected immediately following medium replacement using the PX458-GFP plasmid construct, according to the transfection protocol detailed in the preceding section/Methods. And GFP fluorescence signal was assessed and representative phase-contrast microscopy images were captured at 48 hours post-transfection.

### 6 In vitro digestion of CRISPR system

All in vitro cleavage reaction mixtures were assembled in 50 µL PCR tubes maintained on ice. For standard cleavage reaction, reaction mixtures (5 µL final volume) contained: 0.25 µL Cas9/Cas12/Cas13/dCas9 protein, 0.25 µL positive control substrate (0.5 µL for Cas13), 4 µL nuclease-free water (3.75 µL for Cas13), and 0.5 µL HOMELESS reaction buffer; For reaction with NMN co-incubation, reaction mixtures (5 µL final volume) contained: 0.25 µL Cas9/Cas12/Cas13 protein, 0.25 µL positive control substrate (0.5 µL for Cas13), 4 µL NMN at the indicated concentration (3.75 µL for Cas13), and 0.5 µL HOMELESS reaction buffer; For reaction with NMN-substrate pre-incubation, reaction mixtures (5 µL final volume) contained: 0.25 µL positive control substrate (0.5 µL for Cas13) was pre-mixed with 4 µL NMN at the indicated concentration (3.75 µL for Cas13) and incubated at room temperature for 30 min, subsequently, 0.25 µL Cas9/Cas12/Cas13/dCas9 protein and 0.5 µL HOMELESS reaction buffer were added to the mixture; For reaction with NMN-protein pre-incubation, reaction mixtures (5 µL final volume) contained: 0.25 µL Cas9/Cas12/Cas13/dCas9 protein was pre-mixed with 4 µL NMN at the indicated concentration (3.75 µL for Cas13) and incubated at room temperature for 30 min, subsequently, 0.25 µL positive control substrate (0.5 µL for Cas13) and 0.5 µL HOMELESS reaction buffer were added to the mixture; **For negative control (substrate control),** reaction mixtures (5 µL final volume) contained: 0.25 µL positive control substrate (0.5 µL for Cas13), 4.25 µL nuclease-free water (4 µL for Cas13), and 0.5 µL HOMELESS reaction buffer. Note: This reaction lacks enzyme.Following assembly, all reaction mixtures were thoroughly mixed by pipetting and incubated at 37°C for 30 min in a thermal cycler (PCR machine).

### 7 Visualization detection for DNA by agarose gel electrophoresis

Preparation of Agarose Gel: A total of 1.5 g (1.5%) of agarose was accurately weighted by an electronic balance and transferred into a conical flask. Subsequently, 100 mL of 1× TAE buffer solution was measured by a graduated cylinder and added to the flask. The mixture was thoroughly agitated to ensure homogeneity and then heated in a microwave for 4 min. After heating, the solution was allowed to cool to a temperature range of 50∼60°C. At this point, 10 μL of 10000× nucleic acids dye was incorporated into the solution. The mixture was then poured into a casting tray, and a comb was inserted to form wells. Agarose Gel was left to solidify at room temperature. Sample Loading: 1 μL of 6× DNA Loading Buffer was mixed with 5 μL of CRISPR enzyme digestion products prior to loading into the gel. The voltage was adjusted to 85 V, and the electrophoresis was started for 40 min. Observation: After 40min, the gel was removed and put into an AlphaImager HP gel-imager to observe the results.

### 8 Visualization detection for RNA by SDS-PAGE

Remove the SDS-PAGE gel (GeneScript) and insert it into the SDS-PAGE electrophoresis tank, 1.67 μL of 5× RNA Loading Buffer was mixed with 5 μL of CRISPR enzyme digestion products prior to loading into the gel. The voltage was adjusted to 85 V, and the electrophoresis was started for 60 min. Dyeing: After 60min, the gel was removed and put in a box, add 100ml of 1× nucleic acids dye, and shake it on a shaker for 120min; Observation: put the gel into an AlphaImager HP gel-imager to observe the results.

### 9 qPCR detection for CRISPR enzyme digestion products

According to the specified protocol, the reaction mixture should be prepared as follows: 25 μL of Taq Pro Universal SYBR qPCR Master Mix, 2 μL of each primer, 2 μL of CRISPR enzyme digestion products, and 19 μL of ddH2O, totaling 50 μL. Thermal cycling conditions were as follows: an initial denaturation at 95°C for 5 minutes, followed by 35 cycles of denaturation at 95°C for 30 seconds, annealing at 55°C for 30 seconds, and extension at 72°C for 45 seconds, concluding with a final extension at 72°C for 10 minutes. For each experimental group, add the reagents to the reaction tube, ensure thorough mixing by shaking, and proceed with amplification using the ROUCH fluorescence quantitative PCR system. Upon completion of the reaction, evaluate the results based on the Cq values to draw conclusions.

### 10 EMSA detection for CRISPR system protein-nucleic acids ligation

Remove the Native-PAGE gel (Thermo Scientific) and insert it into the mini gel-tank, 0.67 μL of 4× sample Loading Buffer was mixed with 5 μL of CRISPR d-Cas9 enzyme digestion products prior to loading into the gel. The voltage was adjusted to 85 V, and the electrophoresis was started for 90 min. Dyeing: After 95min, the gel was removed and put in a box, add 100ml of 1× nucleic acids dye, and shake it on a shaker for 15min; Observation: put the gel into an AlphaImager HP gel-imager to observe the results.

### 11 BLI analyses

For DNA and NMN, BLI analyses were performed at 25 °C using the GatorPrime biosensor system (GatorBio) with SA-XT probes (GatorBio,160029). ssDNA with biotin (500 nM) were immobilized onto the anti-Biotin bio sensor for 913 s. The tips were washed with ddH2O for 120 s to obtain a base line reading, then the biosensors were dipped into wells containing the various concentrations of NMN for 240 s, then wash the biosensors were washed for 240 s at ddH2O to dissociate the NMN from the sensor. Data analysis was performed with Graphad Prism 9.0.0 with the One site - specific binding model.

For Cas9 protein and NMN, BLI analyses were performed at 25 °C using the GatorPrime biosensor system (GatorBio) with AR-probes (GatorBio,160008) method. At first, performed buffer exchange on the protein to remove the interference of Tris-HCl from the original buffer, and the bio sensor was put into EDC and sNHS to be activated for 300s. Then Cas9 protein (10μg/ml) were immobilized onto bio sensor for 346 s. After this, the conjugation was quenched by Ethanolamine for 300 s. The tips were washed with ddH2O for 300 s to obtain a base line reading, then the biosensors were dipped into wells containing the various concentrations of NMN for 211 s, then wash the biosensors were washed for 347 s at ddH2O to dissociate the NMN from the sensor. Data analysis was performed with Graphad Prism 9.0.0 with the One site - specific binding model.

### 12 Molecular dynamics

Molecular dynamics (MD) simulations were conducted using the GROMACS 2021.5 software package [28]. The simulation system was established in a confined environment with a temperature of 300 K by the V-rescal thermostat and a pressure of 1 bar by the Parrinello-Rahman barostat. Periodic boundary conditions were applied, with the system centered and the minimum distance between the complex edge and the box edge set to 1.0 nm. The protein and DNA/RNA parameterized by AMBERff14SB force field [29]. Ligand was parameterized by AmberTools [30], and the GAFF force field was applied [31]. The TIP3P model [32] was used for water molecules. After the initial system setup, energy minimization was performed using the steepest descent algorithm with a force tolerance of 500 kJ mol^-1^ nm^-1^. Then system was relaxed for 1000 ps under NVT ensemble and position restraints with a constant of 1000 kJ mol^-1^ nm-^2^ in three directions were performed on heavy atoms of protein/DNA/RNA, followed by a 1000 ps NPT ensemble. Following NVT and NPT equilibration, a 100 ns production MD simulation was performed, with a time step of 2 fs. The LINCS algorithm was performed for constrain bond lengths of hydrogen atoms. Lennard-Jones interactions were calculated within a cutoff of 1.4 nm, and electrostatic interactions beyond 1.4 nm were treated with particle-mesh Ewald (PME) method [33]. After completing all simulations, the gmx module was used to calculate the radius of gyration (Rg), root mean square deviation (RMSD), and root mean square fluctuation (RMSF). The binding free energy (ΔGbind) between the substrate and the system was calculated using the Molecular Mechanics/Generalized Born Surface Area (MM/GBSA) method [34,35].

### 13 Data analysis and result interpretation

All data were repeated three times in each group and averaged for independent samples test, the analysis of variance was used when comparing multiple sets of data with each other (spss20.0 software https://www.ibm.com/spss).

## Authors’ contributions

Tong Wei: Writing – original draft, Investigation, Formal analysis, Conceptualization.

Wei Shen: Writing – review & editing, Methodology, Investigation, Formal analysis.

Wenjun Li: Writing – review & editing, Investigation, Formal analysis.

Yingjie Song: Investigation, Formal analysis.

Yunzhi Fa: Supervision, Resources, Conceptualization.

Yansong Sun: Supervision, Resources, Conceptualization, Funding acquisition.

Hao Li: Writing – review & editing, Supervision, Funding acquisition, Conceptualization.

## Data availability

The data that support the findings of this study are available from the corresponding author, [Yz F], upon reasonable request.

## Ethics approval and consent to participate

Not applicable.

## Consent to publish

Not applicable.

## Acknowledgments

Not applicable

## Animal Ethics

No approval of research ethics committees was required to accomplish the goals of this study because experimental work was conducted with no animal.

## Expand informaintion

**Expand table 1.** Bonding information of Molecular dynamics of NMN and double-stranded DNA.

| name | distance | category | type |
| --- | --- | --- | --- |
| ligand325:H29 - B:DA3:O2P | 1.80859 | Hydrogen Bond | Conventional Hydrogen Bond |
| ligand325:H30 - B:DA3:O2P | 1.64034 | Hydrogen Bond | Conventional Hydrogen Bond |
| ligand327:H29 - A:DG243:O6 | 1.63603 | Hydrogen Bond | Conventional Hydrogen Bond |
| ligand327:H30 - A:DG243:O6 | 3.02506 | Hydrogen Bond | Conventional Hydrogen Bond |
| ligand327:H30 - B:DT20:O4 | 1.94987 | Hydrogen Bond | Conventional Hydrogen Bond |
| ligand327:H37 - A:DG242:O2P | 2.49956 | Hydrogen Bond | Conventional Hydrogen Bond |
| ligand329:H29 - A:DC60:O2 | 1.89729 | Hydrogen Bond | Conventional Hydrogen Bond |
| ligand329:H29 - A:DG61:O4' | 2.73077 | Hydrogen Bond | Conventional Hydrogen Bond |
| ligand329:H30 - B:DA205:N3 | 1.88322 | Hydrogen Bond | Conventional Hydrogen Bond |
| ligand329:H30 - B:DT206:O4' | 2.74725 | Hydrogen Bond | Conventional Hydrogen Bond |
| ligand329:H35 - B:DG207:O1P | 1.85583 | Hydrogen Bond | Conventional Hydrogen Bond |
| ligand331:H29 - A:DC5:O2P | 2.00267 | Hydrogen Bond | Conventional Hydrogen Bond |
| ligand332:H30 - B:DG257:N7 | 2.15358 | Hydrogen Bond | Conventional Hydrogen Bond |
| ligand333:H29 - A:DC206:O1P | 1.59819 | Hydrogen Bond | Conventional Hydrogen Bond |
| ligand333:H36 - B:DG60:N3 | 2.24816 | Hydrogen Bond | Conventional Hydrogen Bond |
| ligand337:H30 - A:DG42:O6 | 1.93902 | Hydrogen Bond | Conventional Hydrogen Bond |
| ligand337:H30 - A:DG43:O6 | 2.4445 | Hydrogen Bond | Conventional Hydrogen Bond |
| ligand337:H35 - A:DA41:O2P | 1.64797 | Hydrogen Bond | Conventional Hydrogen Bond |
| ligand338:H35 - B:DA83:N7 | 1.89093 | Hydrogen Bond | Conventional Hydrogen Bond |
| ligand339:H36 - A:DG193:O4' | 2.69704 | Hydrogen Bond | Conventional Hydrogen Bond |
| ligand339:H37 - A:DT192:O3' | 1.88332 | Hydrogen Bond | Conventional Hydrogen Bond |
| ligand341:H29 - A:DG259:O6 | 2.9544 | Hydrogen Bond | Conventional Hydrogen Bond |
| ligand341:H29 - A:DG259:N7 | 1.94777 | Hydrogen Bond | Conventional Hydrogen Bond |
| ligand341:H30 - A:DG259:O6 | 1.71759 | Hydrogen Bond | Conventional Hydrogen Bond |
| ligand341:H35 - B:DA3:N7 | 1.80258 | Hydrogen Bond | Conventional Hydrogen Bond |
| ligand341:H36 - A:DG258:O6 | 2.03666 | Hydrogen Bond | Conventional Hydrogen Bond |
| ligand342:H29 - B:DC125:O2P | 1.58524 | Hydrogen Bond | Conventional Hydrogen Bond |
| ligand344:H35 - B:DT133:O1P | 1.6568 | Hydrogen Bond | Conventional Hydrogen Bond |
| ligand346:H30 - A:DG52:O6 | 1.92242 | Hydrogen Bond | Conventional Hydrogen Bond |
| ligand346:H35 - A:DG50:O2P | 1.84743 | Hydrogen Bond | Conventional Hydrogen Bond |
| ligand348:H29 - A:DA89:O1P | 1.80624 | Hydrogen Bond | Conventional Hydrogen Bond |
| ligand348:H35 - A:DC88:O1P | 1.76765 | Hydrogen Bond | Conventional Hydrogen Bond |
| ligand353:H29 - A:DC78:O1P | 1.7404 | Hydrogen Bond | Conventional Hydrogen Bond |
| ligand353:H37 - B:DA189:O3' | 2.67836 | Hydrogen Bond | Conventional Hydrogen Bond |
| ligand354:H29 - A:DG52:O2P | 1.88199 | Hydrogen Bond | Conventional Hydrogen Bond |
| ligand354:H30 - A:DG52:O2P | 1.64861 | Hydrogen Bond | Conventional Hydrogen Bond |
| ligand354:H36 - A:DG51:O2P | 1.82877 | Hydrogen Bond | Conventional Hydrogen Bond |
| ligand355:H29 - B:DG61:O2P | 1.58016 | Hydrogen Bond | Conventional Hydrogen Bond |
| ligand355:H35 - B:DG60:O2P | 1.91481 | Hydrogen Bond | Conventional Hydrogen Bond |
| ligand358:H29 - A:DG162:O2P | 2.16539 | Hydrogen Bond | Conventional Hydrogen Bond |
| ligand358:H30 - A:DG162:O2P | 1.66871 | Hydrogen Bond | Conventional Hydrogen Bond |
| ligand358:H37 - B:DG100:N7 | 2.05071 | Hydrogen Bond | Conventional Hydrogen Bond |
| ligand361:H36 - B:DA111:O1P | 2.00803 | Hydrogen Bond | Conventional Hydrogen Bond |
| ligand362:H29 - A:DA17:O2P | 1.54577 | Hydrogen Bond | Conventional Hydrogen Bond |
| ligand362:H30 - A:DA17:N7 | 2.87426 | Hydrogen Bond | Conventional Hydrogen Bond |
| ligand362:H37 - B:DC243:O2P | 1.91256 | Hydrogen Bond | Conventional Hydrogen Bond |
| ligand363:H29 - B:DC228:O2P | 1.82858 | Hydrogen Bond | Conventional Hydrogen Bond |
| ligand363:H35 - B:DT227:O2P | 1.74017 | Hydrogen Bond | Conventional Hydrogen Bond |
| ligand314:H35 - B:DT133:O5' | 1.85329 | Hydrogen Bond | Conventional Hydrogen Bond |
| ligand315:H29 - A:DA165:O2P | 1.89766 | Hydrogen Bond | Conventional Hydrogen Bond |
| ligand315:H30 - A:DA165:O1P | 1.62785 | Hydrogen Bond | Conventional Hydrogen Bond |
| ligand316:H29 - B:DT99:O2 | 2.92193 | Hydrogen Bond | Conventional Hydrogen Bond |
| ligand316:H29 - B:DG100:N3 | 2.02583 | Hydrogen Bond | Conventional Hydrogen Bond |
| ligand316:H30 - B:DT99:O2 | 2.05227 | Hydrogen Bond | Conventional Hydrogen Bond |
| ligand316:H36 - A:DT167:O2 | 2.15037 | Hydrogen Bond | Conventional Hydrogen Bond |
| ligand316:H37 - A:DC168:O4' | 2.16972 | Hydrogen Bond | Conventional Hydrogen Bond |
| ligand318:H29 - B:DC243:O3' | 2.39424 | Hydrogen Bond | Conventional Hydrogen Bond |
| ligand318:H30 - B:DC244:O1P | 2.18845 | Hydrogen Bond | Conventional Hydrogen Bond |
| ligand319:H29 - B:DC6:O2P | 1.66736 | Hydrogen Bond | Conventional Hydrogen Bond |
| ligand319:H30 - B:DC6:O2P | 2.10604 | Hydrogen Bond | Conventional Hydrogen Bond |
| ligand319:H36 - B:DA7:N7 | 2.35991 | Hydrogen Bond | Conventional Hydrogen Bond |
| ligand319:H36 - B:DG8:O6 | 2.62895 | Hydrogen Bond | Conventional Hydrogen Bond |
| ligand321:H29 - B:DC244:O1P | 1.89251 | Hydrogen Bond | Conventional Hydrogen Bond |
| ligand322:H29 - B:DT20:O1P | 1.7663 | Hydrogen Bond | Conventional Hydrogen Bond |
| ligand322:H30 - B:DG19:O3' | 2.63511 | Hydrogen Bond | Conventional Hydrogen Bond |
| ligand322:H37 - A:DC249:O1P | 2.9175 | Hydrogen Bond | Conventional Hydrogen Bond |
| A:DC18:H41 - ligand362:O4 | 2.12904 | Hydrogen Bond | Conventional Hydrogen Bond |
| A:DA34:H61 - ligand363:O9 | 1.87838 | Hydrogen Bond | Conventional Hydrogen Bond |
| A:DC35:H41 - ligand363:O9 | 2.96811 | Hydrogen Bond | Conventional Hydrogen Bond |
| A:DC35:H42 - ligand363:O9 | 2.93123 | Hydrogen Bond | Conventional Hydrogen Bond |
| A:DG54:H22 - ligand288:O3 | 1.83799 | Hydrogen Bond | Conventional Hydrogen Bond |
| A:DG105:H22 - ligand290:O9 | 2.10183 | Hydrogen Bond | Conventional Hydrogen Bond |
| A:DG114:H22 - ligand283:O5 | 1.9956 | Hydrogen Bond | Conventional Hydrogen Bond |
| A:DC163:H41 - ligand358:O5 | 2.51992 | Hydrogen Bond | Conventional Hydrogen Bond |
| A:DC164:H41 - ligand358:O8 | 1.97578 | Hydrogen Bond | Conventional Hydrogen Bond |
| A:DC170:H41 - ligand307:O5 | 2.98767 | Hydrogen Bond | Conventional Hydrogen Bond |
| A:DT192:H3 - ligand339:O4 | 2.56271 | Hydrogen Bond | Conventional Hydrogen Bond |
| B:DA3:H61 - ligand341:O7 | 2.3692 | Hydrogen Bond | Conventional Hydrogen Bond |
| B:DC62:H41 - ligand265:O4 | 2.29133 | Hydrogen Bond | Conventional Hydrogen Bond |
| B:DA84:H61 - ligand338:O5 | 2.164 | Hydrogen Bond | Conventional Hydrogen Bond |
| B:DA126:H61 - ligand342:O4 | 2.07446 | Hydrogen Bond | Conventional Hydrogen Bond |
| B:DG149:H22 - ligand283:O3 | 1.80236 | Hydrogen Bond | Conventional Hydrogen Bond |
| B:DG196:H22 - ligand357:O4 | 1.84686 | Hydrogen Bond | Conventional Hydrogen Bond |
| B:DG204:H22 - ligand284:O4 | 2.1487 | Hydrogen Bond | Conventional Hydrogen Bond |
| B:DC238:H41 - ligand278:O7 | 2.12462 | Hydrogen Bond | Conventional Hydrogen Bond |
| B:DG257:H22 - ligand291:O3 | 2.25579 | Hydrogen Bond | Conventional Hydrogen Bond |
| B:DG257:H22 - ligand291:O4 | 2.17039 | Hydrogen Bond | Conventional Hydrogen Bond |
| ligand265:H29 - B:DG61:O2P | 1.79048 | Hydrogen Bond | Conventional Hydrogen Bond |
| ligand265:H30 - B:DG61:N7 | 2.55204 | Hydrogen Bond | Conventional Hydrogen Bond |
| ligand265:H37 - A:DA199:O2P | 1.7912 | Hydrogen Bond | Conventional Hydrogen Bond |
| ligand268:H35 - B:DT217:O2P | 1.82225 | Hydrogen Bond | Conventional Hydrogen Bond |
| ligand270:H29 - A:DC7:O2P | 1.71155 | Hydrogen Bond | Conventional Hydrogen Bond |
| ligand270:H30 - A:DA8:O2P | 1.52643 | Hydrogen Bond | Conventional Hydrogen Bond |
| ligand272:H35 - A:DG103:O2P | 1.75585 | Hydrogen Bond | Conventional Hydrogen Bond |
| ligand272:H36 - B:DT163:O2 | 1.76819 | Hydrogen Bond | Conventional Hydrogen Bond |
| ligand272:H37 - A:DC102:O2 | 1.98409 | Hydrogen Bond | Conventional Hydrogen Bond |
| ligand274:H37 - A:DC78:O1P | 1.95267 | Hydrogen Bond | Conventional Hydrogen Bond |
| ligand278:H29 - B:DC238:O2P | 1.73254 | Hydrogen Bond | Conventional Hydrogen Bond |
| ligand278:H35 - B:DG237:N7 | 2.09347 | Hydrogen Bond | Conventional Hydrogen Bond |
| ligand280:H29 - B:DA250:O2P | 1.74997 | Hydrogen Bond | Conventional Hydrogen Bond |
| ligand280:H30 - B:DA250:O2P | 1.60866 | Hydrogen Bond | Conventional Hydrogen Bond |
| ligand280:H36 - B:DT251:O2P | 1.63297 | Hydrogen Bond | Conventional Hydrogen Bond |
| ligand281:H29 - B:DG249:O2P | 2.9828 | Hydrogen Bond | Conventional Hydrogen Bond |
| ligand281:H30 - B:DG249:O2P | 1.67771 | Hydrogen Bond | Conventional Hydrogen Bond |
| ligand281:H36 - B:DG249:O1P | 1.84556 | Hydrogen Bond | Conventional Hydrogen Bond |
| ligand282:H36 - B:DC215:O2 | 1.91458 | Hydrogen Bond | Conventional Hydrogen Bond |
| ligand283:H35 - A:DC115:O2 | 1.74462 | Hydrogen Bond | Conventional Hydrogen Bond |
| ligand285:H30 - A:DG189:O6 | 1.8348 | Hydrogen Bond | Conventional Hydrogen Bond |
| ligand287:H30 - B:DC132:O1P | 1.74809 | Hydrogen Bond | Conventional Hydrogen Bond |
| ligand287:H35 - A:DC135:O1P | 1.58424 | Hydrogen Bond | Conventional Hydrogen Bond |
| ligand288:H30 - A:DG55:O4' | 2.03995 | Hydrogen Bond | Conventional Hydrogen Bond |
| ligand291:H29 - B:DG257:N3 | 2.57901 | Hydrogen Bond | Conventional Hydrogen Bond |
| ligand291:H30 - A:DA8:N3 | 1.7057 | Hydrogen Bond | Conventional Hydrogen Bond |
| ligand293:H29 - A:DG20:O6 | 1.80483 | Hydrogen Bond | Conventional Hydrogen Bond |
| ligand293:H30 - A:DG21:O6 | 1.69685 | Hydrogen Bond | Conventional Hydrogen Bond |
| ligand293:H36 - A:DG21:N7 | 2.62059 | Hydrogen Bond | Conventional Hydrogen Bond |
| ligand295:H36 - A:DC33:O2 | 1.77862 | Hydrogen Bond | Conventional Hydrogen Bond |
| ligand295:H37 - A:DC33:O4' | 2.79202 | Hydrogen Bond | Conventional Hydrogen Bond |
| ligand296:H29 - A:DC10:O5' | 1.87774 | Hydrogen Bond | Conventional Hydrogen Bond |
| ligand299:H29 - B:DT248:O2 | 2.13059 | Hydrogen Bond | Conventional Hydrogen Bond |
| ligand299:H29 - B:DG249:O4' | 2.80109 | Hydrogen Bond | Conventional Hydrogen Bond |
| ligand299:H35 - B:DG249:O1P | 2.12756 | Hydrogen Bond | Conventional Hydrogen Bond |
| ligand300:H30 - A:DT197:O2P | 1.66069 | Hydrogen Bond | Conventional Hydrogen Bond |
| ligand300:H35 - B:DC62:O2P | 1.88383 | Hydrogen Bond | Conventional Hydrogen Bond |
| ligand300:H37 - B:DG61:O5' | 2.05567 | Hydrogen Bond | Conventional Hydrogen Bond |
| ligand301:H29 - B:DC244:O1P | 1.57016 | Hydrogen Bond | Conventional Hydrogen Bond |
| ligand301:H30 - B:DC244:O2P | 1.71038 | Hydrogen Bond | Conventional Hydrogen Bond |
| ligand302:H29 - A:DA16:O2P | 1.81063 | Hydrogen Bond | Conventional Hydrogen Bond |
| ligand302:H35 - A:DC15:O2P | 1.77313 | Hydrogen Bond | Conventional Hydrogen Bond |
| ligand302:H37 - B:DT247:O4 | 2.24219 | Hydrogen Bond | Conventional Hydrogen Bond |
| ligand303:H30 - B:DG134:O2P | 1.70029 | Hydrogen Bond | Conventional Hydrogen Bond |
| ligand304:H30 - A:DA38:O1P | 1.77389 | Hydrogen Bond | Conventional Hydrogen Bond |
| ligand306:H29 - A:DA195:N7 | 1.82266 | Hydrogen Bond | Conventional Hydrogen Bond |
| ligand306:H35 - B:DG66:N7 | 2.34721 | Hydrogen Bond | Conventional Hydrogen Bond |
| ligand307:H29 - B:DG95:O6 | 1.82327 | Hydrogen Bond | Conventional Hydrogen Bond |
| ligand307:H37 - B:DC92:O2P | 1.92065 | Hydrogen Bond | Conventional Hydrogen Bond |
| ligand310:H29 - A:DG161:O1P | 1.65729 | Hydrogen Bond | Conventional Hydrogen Bond |
| ligand310:H35 - B:DG107:O1P | 2.11367 | Hydrogen Bond | Conventional Hydrogen Bond |
| ligand310:H36 - A:DC159:O2 | 2.02453 | Hydrogen Bond | Conventional Hydrogen Bond |
| ligand310:H36 - A:DT160:O4' | 3.08558 | Hydrogen Bond | Conventional Hydrogen Bond |
| ligand311:H36 - B:DG166:O1P | 2.00472 | Hydrogen Bond | Conventional Hydrogen Bond |
| ligand312:H30 - A:DA53:O2P | 1.61759 | Hydrogen Bond | Conventional Hydrogen Bond |
| ligand312:H37 - A:DG54:N7 | 2.50675 | Hydrogen Bond | Conventional Hydrogen Bond |
| ligand313:H35 - A:DT9:O2P | 1.81417 | Hydrogen Bond | Conventional Hydrogen Bond |
| ligand327:H23 - A:DG243:N7 | 2.98843 | Hydrogen Bond | Carbon Hydrogen Bond |
| ligand327:H24 - A:DG243:O6 | 2.8217 | Hydrogen Bond | Carbon Hydrogen Bond |
| ligand327:H24 - A:DG243:N7 | 2.61247 | Hydrogen Bond | Carbon Hydrogen Bond |
| ligand327:H27 - A:DG242:N7 | 2.57507 | Hydrogen Bond | Carbon Hydrogen Bond |
| ligand327:H31 - A:DG242:N7 | 3.01498 | Hydrogen Bond | Carbon Hydrogen Bond |
| ligand328:H23 - B:DC102:O3' | 2.66582 | Hydrogen Bond | Carbon Hydrogen Bond |
| ligand328:H26 - A:DC164:O3' | 2.38703 | Hydrogen Bond | Carbon Hydrogen Bond |
| ligand328:H26 - A:DA165:O1P | 2.51581 | Hydrogen Bond | Carbon Hydrogen Bond |
| ligand328:H28 - B:DC103:O1P | 2.52565 | Hydrogen Bond | Carbon Hydrogen Bond |
| ligand329:H23 - A:DC60:O2 | 2.59257 | Hydrogen Bond | Carbon Hydrogen Bond |
| ligand329:H24 - A:DC60:O2 | 2.34608 | Hydrogen Bond | Carbon Hydrogen Bond |
| ligand329:H26 - B:DT206:O4' | 2.87906 | Hydrogen Bond | Carbon Hydrogen Bond |
| ligand329:H32 - A:DT59:O2 | 2.46311 | Hydrogen Bond | Carbon Hydrogen Bond |
| ligand329:H32 - B:DT206:O2 | 2.76846 | Hydrogen Bond | Carbon Hydrogen Bond |
| ligand332:H23 - B:DG257:O6 | 2.23325 | Hydrogen Bond | Carbon Hydrogen Bond |
| ligand332:H28 - B:DG257:O6 | 2.22182 | Hydrogen Bond | Carbon Hydrogen Bond |
| ligand333:H25 - A:DC206:O1P | 2.95491 | Hydrogen Bond | Carbon Hydrogen Bond |
| ligand333:H26 - A:DC205:O3' | 2.24308 | Hydrogen Bond | Carbon Hydrogen Bond |
| ligand333:H32 - A:DC205:O4' | 2.68159 | Hydrogen Bond | Carbon Hydrogen Bond |
| ligand337:H23 - A:DG42:O6 | 2.62013 | Hydrogen Bond | Carbon Hydrogen Bond |
| ligand337:H24 - A:DG42:N7 | 2.61245 | Hydrogen Bond | Carbon Hydrogen Bond |
| ligand337:H28 - A:DG42:N7 | 2.88115 | Hydrogen Bond | Carbon Hydrogen Bond |
| ligand339:H26 - B:DG73:N7 | 2.61255 | Hydrogen Bond | Carbon Hydrogen Bond |
| ligand339:H31 - A:DT192:O2 | 2.69316 | Hydrogen Bond | Carbon Hydrogen Bond |
| ligand339:H32 - B:DA72:O3' | 2.5102 | Hydrogen Bond | Carbon Hydrogen Bond |
| ligand341:H23 - A:DG259:O6 | 2.90534 | Hydrogen Bond | Carbon Hydrogen Bond |
| ligand341:H23 - B:DG4:O6 | 2.6841 | Hydrogen Bond | Carbon Hydrogen Bond |
| ligand341:H24 - B:DG4:O6 | 3.04229 | Hydrogen Bond | Carbon Hydrogen Bond |
| ligand341:H27 - B:DG4:O6 | 2.58103 | Hydrogen Bond | Carbon Hydrogen Bond |
| ligand341:H28 - B:DG4:O6 | 2.52719 | Hydrogen Bond | Carbon Hydrogen Bond |
| ligand341:H31 - A:DG258:N7 | 3.0691 | Hydrogen Bond | Carbon Hydrogen Bond |
| ligand342:H23 - B:DA126:N7 | 2.56314 | Hydrogen Bond | Carbon Hydrogen Bond |
| ligand342:H25 - B:DC125:O2P | 2.42526 | Hydrogen Bond | Carbon Hydrogen Bond |
| ligand346:H24 - A:DG51:O6 | 2.4575 | Hydrogen Bond | Carbon Hydrogen Bond |
| ligand346:H24 - A:DG52:O6 | 2.8242 | Hydrogen Bond | Carbon Hydrogen Bond |
| ligand346:H32 - B:DT211:O4 | 2.56332 | Hydrogen Bond | Carbon Hydrogen Bond |
| ligand347:H23 - B:DG249:O5' | 2.26775 | Hydrogen Bond | Carbon Hydrogen Bond |
| ligand347:H24 - B:DG249:O2P | 2.66565 | Hydrogen Bond | Carbon Hydrogen Bond |
| ligand347:H27 - B:DG249:O2P | 2.60348 | Hydrogen Bond | Carbon Hydrogen Bond |
| ligand354:H23 - A:DG52:O2P | 2.63566 | Hydrogen Bond | Carbon Hydrogen Bond |
| ligand354:H31 - A:DG51:O2P | 2.51843 | Hydrogen Bond | Carbon Hydrogen Bond |
| ligand355:H25 - B:DG61:O2P | 3.0953 | Hydrogen Bond | Carbon Hydrogen Bond |
| ligand357:H26 - B:DG198:O4' | 2.63674 | Hydrogen Bond | Carbon Hydrogen Bond |
| ligand357:H32 - B:DT197:O2 | 2.63945 | Hydrogen Bond | Carbon Hydrogen Bond |
| ligand358:H23 - A:DG162:O2P | 2.81306 | Hydrogen Bond | Carbon Hydrogen Bond |
| ligand361:H31 - B:DA111:O1P | 2.73148 | Hydrogen Bond | Carbon Hydrogen Bond |
| ligand362:H25 - A:DA17:O2P | 3.05828 | Hydrogen Bond | Carbon Hydrogen Bond |
| ligand362:H32 - B:DG245:O6 | 2.83106 | Hydrogen Bond | Carbon Hydrogen Bond |
| ligand363:H32 - B:DG229:N7 | 2.63443 | Hydrogen Bond | Carbon Hydrogen Bond |
| ligand314:H23 - B:DG134:O6 | 2.79718 | Hydrogen Bond | Carbon Hydrogen Bond |
| ligand314:H23 - B:DG134:N7 | 2.46329 | Hydrogen Bond | Carbon Hydrogen Bond |
| ligand314:H28 - B:DG134:N7 | 2.44542 | Hydrogen Bond | Carbon Hydrogen Bond |
| ligand316:H23 - B:DG100:O4' | 2.48373 | Hydrogen Bond | Carbon Hydrogen Bond |
| ligand316:H26 - A:DC166:O2 | 2.2802 | Hydrogen Bond | Carbon Hydrogen Bond |
| ligand316:H31 - A:DC166:O2 | 2.46953 | Hydrogen Bond | Carbon Hydrogen Bond |
| ligand316:H31 - A:DT167:O4' | 2.60373 | Hydrogen Bond | Carbon Hydrogen Bond |
| ligand318:H23 - B:DC244:O1P | 2.52438 | Hydrogen Bond | Carbon Hydrogen Bond |
| ligand318:H24 - B:DC244:O1P | 3.04862 | Hydrogen Bond | Carbon Hydrogen Bond |
| ligand319:H23 - B:DC6:O2P | 2.69299 | Hydrogen Bond | Carbon Hydrogen Bond |
| ligand319:H27 - B:DA7:O2P | 2.83868 | Hydrogen Bond | Carbon Hydrogen Bond |
| ligand321:H23 - B:DC244:O1P | 2.38155 | Hydrogen Bond | Carbon Hydrogen Bond |
| ligand321:H32 - B:DG245:O1P | 2.35519 | Hydrogen Bond | Carbon Hydrogen Bond |
| ligand322:H23 - B:DT20:O1P | 3.01354 | Hydrogen Bond | Carbon Hydrogen Bond |
| ligand323:H32 - B:DA189:N3 | 2.67531 | Hydrogen Bond | Carbon Hydrogen Bond |
| A:DC5:H6 - ligand332:O6 | 3.0801 | Hydrogen Bond | Carbon Hydrogen Bond |
| A:DC15:H6 - ligand302:O2 | 2.8632 | Hydrogen Bond | Carbon Hydrogen Bond |
| A:DA16:H8 - ligand302:O3 | 2.54313 | Hydrogen Bond | Carbon Hydrogen Bond |
| A:DC18:H4' - ligand299:O7 | 2.53831 | Hydrogen Bond | Carbon Hydrogen Bond |
| A:DG24:H5'1 - ligand318:O2 | 2.54515 | Hydrogen Bond | Carbon Hydrogen Bond |
| A:DA41:H8 - ligand337:O5 | 2.56657 | Hydrogen Bond | Carbon Hydrogen Bond |
| A:DG51:H4' - ligand282:O6 | 2.96203 | Hydrogen Bond | Carbon Hydrogen Bond |
| A:DG51:H4' - ligand282:O9 | 2.82868 | Hydrogen Bond | Carbon Hydrogen Bond |
| A:DG51:H5'2 - ligand282:O2 | 2.73925 | Hydrogen Bond | Carbon Hydrogen Bond |
| A:DG51:H5'2 - ligand282:O6 | 2.88834 | Hydrogen Bond | Carbon Hydrogen Bond |
| A:DG54:H1' - ligand288:O4 | 2.30413 | Hydrogen Bond | Carbon Hydrogen Bond |
| A:DC68:H4' - ligand357:O9 | 2.7554 | Hydrogen Bond | Carbon Hydrogen Bond |
| A:DA77:H4' - ligand323:O2 | 2.57342 | Hydrogen Bond | Carbon Hydrogen Bond |
| A:DC88:H5'2 - ligand348:O5 | 2.48889 | Hydrogen Bond | Carbon Hydrogen Bond |
| A:DC102:H1' - ligand272:O9 | 2.30109 | Hydrogen Bond | Carbon Hydrogen Bond |
| A:DC102:H4' - ligand272:O2 | 2.57144 | Hydrogen Bond | Carbon Hydrogen Bond |
| A:DC102:H5'2 - ligand272:O5 | 3.08243 | Hydrogen Bond | Carbon Hydrogen Bond |
| A:DG107:H4' - ligand290:O4 | 2.84352 | Hydrogen Bond | Carbon Hydrogen Bond |
| A:DT116:H4' - ligand283:O2 | 2.81311 | Hydrogen Bond | Carbon Hydrogen Bond |
| A:DT116:H5'1 - ligand283:O6 | 2.64913 | Hydrogen Bond | Carbon Hydrogen Bond |
| A:DC157:H4' - ligand361:O5 | 2.90412 | Hydrogen Bond | Carbon Hydrogen Bond |
| A:DC157:H5'2 - ligand361:O2 | 3.04076 | Hydrogen Bond | Carbon Hydrogen Bond |
| A:DC157:H5'2 - ligand361:O5 | 2.43345 | Hydrogen Bond | Carbon Hydrogen Bond |
| A:DC164:H4' - ligand328:O3 | 2.61339 | Hydrogen Bond | Carbon Hydrogen Bond |
| A:DA165:H4' - ligand328:O9 | 2.65255 | Hydrogen Bond | Carbon Hydrogen Bond |
| A:DA165:H5'1 - ligand328:O9 | 2.75949 | Hydrogen Bond | Carbon Hydrogen Bond |
| A:DC168:H4' - ligand316:O9 | 2.41752 | Hydrogen Bond | Carbon Hydrogen Bond |
| A:DC168:H5'1 - ligand316:O9 | 2.4681 | Hydrogen Bond | Carbon Hydrogen Bond |
| A:DC198:H6 - ligand300:O3 | 2.65437 | Hydrogen Bond | Carbon Hydrogen Bond |
| A:DA199:H8 - ligand265:O9 | 2.77325 | Hydrogen Bond | Carbon Hydrogen Bond |
| A:DC206:H5'2 - ligand333:O2 | 2.79739 | Hydrogen Bond | Carbon Hydrogen Bond |
| A:DC206:H5'2 - ligand333:O7 | 2.50633 | Hydrogen Bond | Carbon Hydrogen Bond |
| A:DC227:H4' - ligand264:O3 | 2.4264 | Hydrogen Bond | Carbon Hydrogen Bond |
| A:DG248:H4' - ligand322:O2 | 2.72767 | Hydrogen Bond | Carbon Hydrogen Bond |
| B:DC6:H6 - ligand319:O4 | 2.3234 | Hydrogen Bond | Carbon Hydrogen Bond |
| B:DT18:H1' - ligand322:O4 | 2.45304 | Hydrogen Bond | Carbon Hydrogen Bond |
| B:DC39:H1' - ligand264:O4 | 2.79525 | Hydrogen Bond | Carbon Hydrogen Bond |
| B:DG60:H3' - ligand355:O7 | 2.49884 | Hydrogen Bond | Carbon Hydrogen Bond |
| B:DG61:H8 - ligand265:O3 | 2.71358 | Hydrogen Bond | Carbon Hydrogen Bond |
| B:DG61:H5'1 - ligand333:O9 | 2.67054 | Hydrogen Bond | Carbon Hydrogen Bond |
| B:DC62:H4' - ligand333:O4 | 2.89344 | Hydrogen Bond | Carbon Hydrogen Bond |
| B:DC62:H5'1 - ligand333:O4 | 3.08406 | Hydrogen Bond | Carbon Hydrogen Bond |
| B:DC62:H5'2 - ligand333:O4 | 2.85072 | Hydrogen Bond | Carbon Hydrogen Bond |
| B:DT99:H1' - ligand316:O4 | 2.56875 | Hydrogen Bond | Carbon Hydrogen Bond |
| B:DG107:H4' - ligand310:O9 | 2.68749 | Hydrogen Bond | Carbon Hydrogen Bond |
| B:DG107:H5'1 - ligand310:O9 | 2.89883 | Hydrogen Bond | Carbon Hydrogen Bond |
| B:DT133:H6 - ligand314:O5 | 2.34357 | Hydrogen Bond | Carbon Hydrogen Bond |
| B:DG165:H4' - ligand311:O4 | 2.26438 | Hydrogen Bond | Carbon Hydrogen Bond |
| B:DA189:H5'2 - ligand353:O9 | 2.62787 | Hydrogen Bond | Carbon Hydrogen Bond |
| B:DT197:H4' - ligand357:O3 | 2.55457 | Hydrogen Bond | Carbon Hydrogen Bond |
| B:DG198:H4' - ligand357:O2 | 2.54026 | Hydrogen Bond | Carbon Hydrogen Bond |
| B:DG198:H4' - ligand357:O7 | 2.76009 | Hydrogen Bond | Carbon Hydrogen Bond |
| B:DA199:H4' - ligand357:O9 | 2.70083 | Hydrogen Bond | Carbon Hydrogen Bond |
| B:DG207:H5'1 - ligand329:O2 | 2.42596 | Hydrogen Bond | Carbon Hydrogen Bond |
| B:DG207:H5'1 - ligand329:O5 | 3.05555 | Hydrogen Bond | Carbon Hydrogen Bond |
| B:DT211:H5'1 - ligand288:O7 | 2.91206 | Hydrogen Bond | Carbon Hydrogen Bond |
| B:DC212:H5'1 - ligand288:O2 | 2.79632 | Hydrogen Bond | Carbon Hydrogen Bond |
| B:DC213:H5'1 - ligand288:O9 | 2.66109 | Hydrogen Bond | Carbon Hydrogen Bond |
| B:DT217:H3' - ligand268:O8 | 2.96759 | Hydrogen Bond | Carbon Hydrogen Bond |
| B:DC228:H6 - ligand363:O3 | 2.5137 | Hydrogen Bond | Carbon Hydrogen Bond |
| B:DG237:H8 - ligand278:O5 | 2.31089 | Hydrogen Bond | Carbon Hydrogen Bond |
| B:DC238:H6 - ligand278:O3 | 2.68255 | Hydrogen Bond | Carbon Hydrogen Bond |
| B:DC244:H4' - ligand321:O4 | 2.90388 | Hydrogen Bond | Carbon Hydrogen Bond |
| B:DC244:H5'2 - ligand321:O4 | 2.37019 | Hydrogen Bond | Carbon Hydrogen Bond |
| B:DT248:H4' - ligand299:O5 | 3.02615 | Hydrogen Bond | Carbon Hydrogen Bond |
| B:DG252:H8 - ligand280:O9 | 2.70972 | Hydrogen Bond | Carbon Hydrogen Bond |
| B:DG258:H5'2 - ligand291:O6 | 2.43041 | Hydrogen Bond | Carbon Hydrogen Bond |
| ligand264:H23 - B:DC40:O1P | 2.56675 | Hydrogen Bond | Carbon Hydrogen Bond |
| ligand264:H24 - B:DC39:O3' | 3.02848 | Hydrogen Bond | Carbon Hydrogen Bond |
| ligand264:H26 - A:DC227:O3' | 2.13687 | Hydrogen Bond | Carbon Hydrogen Bond |
| ligand264:H27 - B:DC39:O3' | 2.64456 | Hydrogen Bond | Carbon Hydrogen Bond |
| ligand264:H28 - B:DC40:O1P | 2.35975 | Hydrogen Bond | Carbon Hydrogen Bond |
| ligand264:H32 - A:DC227:O3' | 3.04345 | Hydrogen Bond | Carbon Hydrogen Bond |
| ligand272:H27 - A:DC102:O3' | 2.93329 | Hydrogen Bond | Carbon Hydrogen Bond |
| ligand272:H32 - B:DG165:O4' | 2.42417 | Hydrogen Bond | Carbon Hydrogen Bond |
| ligand280:H23 - B:DA250:O5' | 2.42971 | Hydrogen Bond | Carbon Hydrogen Bond |
| ligand280:H27 - B:DT251:O2P | 2.67286 | Hydrogen Bond | Carbon Hydrogen Bond |
| ligand280:H31 - B:DT251:O2P | 2.70387 | Hydrogen Bond | Carbon Hydrogen Bond |
| ligand281:H26 - B:DG249:O2P | 2.81947 | Hydrogen Bond | Carbon Hydrogen Bond |
| ligand281:H31 - B:DG249:O1P | 2.32323 | Hydrogen Bond | Carbon Hydrogen Bond |
| ligand283:H23 - B:DC150:O2 | 2.71046 | Hydrogen Bond | Carbon Hydrogen Bond |
| ligand283:H24 - B:DA151:O4' | 2.74423 | Hydrogen Bond | Carbon Hydrogen Bond |
| ligand283:H25 - A:DC115:O2 | 2.69422 | Hydrogen Bond | Carbon Hydrogen Bond |
| ligand283:H27 - B:DA151:O4' | 2.77802 | Hydrogen Bond | Carbon Hydrogen Bond |
| ligand283:H28 - B:DC150:O2 | 2.45088 | Hydrogen Bond | Carbon Hydrogen Bond |
| ligand283:H31 - B:DA151:O4' | 2.91235 | Hydrogen Bond | Carbon Hydrogen Bond |
| ligand284:H26 - A:DA62:O4' | 2.64414 | Hydrogen Bond | Carbon Hydrogen Bond |
| ligand284:H27 - B:DA205:O3' | 3.03873 | Hydrogen Bond | Carbon Hydrogen Bond |
| ligand284:H32 - B:DG204:N3 | 2.9067 | Hydrogen Bond | Carbon Hydrogen Bond |
| ligand285:H23 - A:DA187:N7 | 2.34147 | Hydrogen Bond | Carbon Hydrogen Bond |
| ligand285:H24 - A:DG189:N7 | 2.77939 | Hydrogen Bond | Carbon Hydrogen Bond |
| ligand285:H27 - A:DG188:N7 | 2.71002 | Hydrogen Bond | Carbon Hydrogen Bond |
| ligand285:H28 - A:DG188:N7 | 2.35085 | Hydrogen Bond | Carbon Hydrogen Bond |
| ligand285:H32 - A:DG189:O6 | 2.38908 | Hydrogen Bond | Carbon Hydrogen Bond |
| ligand285:H32 - B:DG74:O6 | 3.07104 | Hydrogen Bond | Carbon Hydrogen Bond |
| ligand286:H26 - B:DG139:N7 | 2.27037 | Hydrogen Bond | Carbon Hydrogen Bond |
| ligand286:H32 - B:DG139:O6 | 2.77433 | Hydrogen Bond | Carbon Hydrogen Bond |
| ligand286:H32 - B:DG139:N7 | 3.08446 | Hydrogen Bond | Carbon Hydrogen Bond |
| ligand287:H24 - B:DC132:O1P | 2.93963 | Hydrogen Bond | Carbon Hydrogen Bond |
| ligand288:H23 - A:DG55:O4' | 2.63169 | Hydrogen Bond | Carbon Hydrogen Bond |
| ligand288:H26 - B:DT211:O2 | 2.50452 | Hydrogen Bond | Carbon Hydrogen Bond |
| ligand290:H24 - A:DG107:O5' | 2.88978 | Hydrogen Bond | Carbon Hydrogen Bond |
| ligand290:H27 - A:DG107:O1P | 2.68119 | Hydrogen Bond | Carbon Hydrogen Bond |
| ligand290:H31 - A:DG107:O1P | 3.08809 | Hydrogen Bond | Carbon Hydrogen Bond |
| ligand291:H23 - A:DT9:O4' | 3.06571 | Hydrogen Bond | Carbon Hydrogen Bond |
| ligand291:H26 - B:DG258:O4' | 2.36738 | Hydrogen Bond | Carbon Hydrogen Bond |
| ligand291:H27 - A:DT9:O3' | 3.0248 | Hydrogen Bond | Carbon Hydrogen Bond |
| ligand293:H23 - A:DG20:N7 | 2.78672 | Hydrogen Bond | Carbon Hydrogen Bond |
| ligand293:H24 - A:DG20:O6 | 2.73388 | Hydrogen Bond | Carbon Hydrogen Bond |
| ligand293:H24 - A:DG20:N7 | 2.73002 | Hydrogen Bond | Carbon Hydrogen Bond |
| ligand293:H24 - A:DG21:O6 | 2.87252 | Hydrogen Bond | Carbon Hydrogen Bond |
| ligand293:H31 - A:DG21:N7 | 2.57499 | Hydrogen Bond | Carbon Hydrogen Bond |
| ligand296:H23 - A:DC10:O2P | 2.83729 | Hydrogen Bond | Carbon Hydrogen Bond |
| ligand296:H26 - A:DA11:O2P | 2.69987 | Hydrogen Bond | Carbon Hydrogen Bond |
| ligand299:H23 - B:DT248:O2 | 2.49636 | Hydrogen Bond | Carbon Hydrogen Bond |
| ligand299:H24 - A:DA17:N3 | 3.02354 | Hydrogen Bond | Carbon Hydrogen Bond |
| ligand299:H25 - B:DT248:O2 | 2.90252 | Hydrogen Bond | Carbon Hydrogen Bond |
| ligand299:H25 - B:DG249:O4' | 2.62513 | Hydrogen Bond | Carbon Hydrogen Bond |
| ligand299:H28 - B:DT248:O2 | 2.91155 | Hydrogen Bond | Carbon Hydrogen Bond |
| ligand300:H24 - A:DT197:O2P | 3.01015 | Hydrogen Bond | Carbon Hydrogen Bond |
| ligand302:H31 - A:DA16:N7 | 2.34455 | Hydrogen Bond | Carbon Hydrogen Bond |
| ligand304:H24 - A:DA38:O1P | 2.8382 | Hydrogen Bond | Carbon Hydrogen Bond |
| ligand304:H24 - A:DA38:O2P | 2.62833 | Hydrogen Bond | Carbon Hydrogen Bond |
| ligand306:H23 - A:DG196:O6 | 2.41512 | Hydrogen Bond | Carbon Hydrogen Bond |
| ligand306:H28 - A:DG196:O6 | 2.64934 | Hydrogen Bond | Carbon Hydrogen Bond |
| ligand307:H23 - B:DG94:O6 | 2.40915 | Hydrogen Bond | Carbon Hydrogen Bond |
| ligand307:H23 - B:DG94:N7 | 2.55229 | Hydrogen Bond | Carbon Hydrogen Bond |
| ligand307:H28 - B:DG94:O6 | 2.47457 | Hydrogen Bond | Carbon Hydrogen Bond |
| ligand310:H24 - A:DT160:O3' | 2.73388 | Hydrogen Bond | Carbon Hydrogen Bond |
| ligand310:H24 - A:DG161:O1P | 2.76854 | Hydrogen Bond | Carbon Hydrogen Bond |
| ligand311:H24 - B:DG165:O3' | 2.75918 | Hydrogen Bond | Carbon Hydrogen Bond |
| ligand311:H31 - B:DG166:O1P | 2.51891 | Hydrogen Bond | Carbon Hydrogen Bond |
| ligand312:H24 - A:DA53:O2P | 2.72232 | Hydrogen Bond | Carbon Hydrogen Bond |
| ligand316:H29 - B:DG100 | 2.84963 | Hydrogen Bond | Pi-Donor Hydrogen Bond |
| A:DG51:H5'1 - ligand282 | 2.62921 | Hydrophobic | Pi-Sigma |
| A:DC228:H5'1 - ligand264 | 2.63296 | Hydrophobic | Pi-Sigma |
| B:DC132:H5'2 - ligand287 | 2.7935 | Hydrophobic | Pi-Sigma |
| ligand329 - B:DG207 | 5.29787 | Hydrophobic | Pi-Pi Stacked |
| ligand339 - B:DG73 | 4.95163 | Hydrophobic | Pi-Pi Stacked |
| ligand357 - A:DC68 | 5.73695 | Hydrophobic | Pi-Pi Stacked |
| ligand362 - B:DC244 | 4.81422 | Hydrophobic | Pi-Pi Stacked |
| A:DC15 - ligand302 | 5.20144 | Hydrophobic | Pi-Pi Stacked |
| B:DG73 - ligand339 | 4.15027 | Hydrophobic | Pi-Pi Stacked |
| B:DG198 - ligand357 | 5.11369 | Hydrophobic | Pi-Pi Stacked |
| B:DA216 - ligand282 | 5.47157 | Hydrophobic | Pi-Pi Stacked |
| B:DA216 - ligand282 | 5.10345 | Hydrophobic | Pi-Pi Stacked |
| ligand346 - A:DG52 | 5.34435 | Hydrophobic | Pi-Pi T-shaped |
| A:DG51 - ligand346 | 5.35376 | Hydrophobic | Pi-Pi T-shaped |
| A:DG52 - ligand346 | 4.42344 | Hydrophobic | Pi-Pi T-shaped |
| A:DG189 - ligand285 | 5.15056 | Hydrophobic | Pi-Pi T-shaped |
| A:DG189 - ligand285 | 4.22829 | Hydrophobic | Pi-Pi T-shaped |
| A:DG242 - ligand327 | 5.00246 | Hydrophobic | Pi-Pi T-shaped |
| A:DG243 - ligand327 | 4.87967 | Hydrophobic | Pi-Pi T-shaped |
| B:DA138 - ligand286 | 5.25411 | Hydrophobic | Pi-Pi T-shaped |
| B:DA250 - ligand320 | 5.14477 | Hydrophobic | Pi-Alkyl |
| A: A chain of dsDNA; B: B chain of dsDNA |  |  |  |

**Expand table 2.**
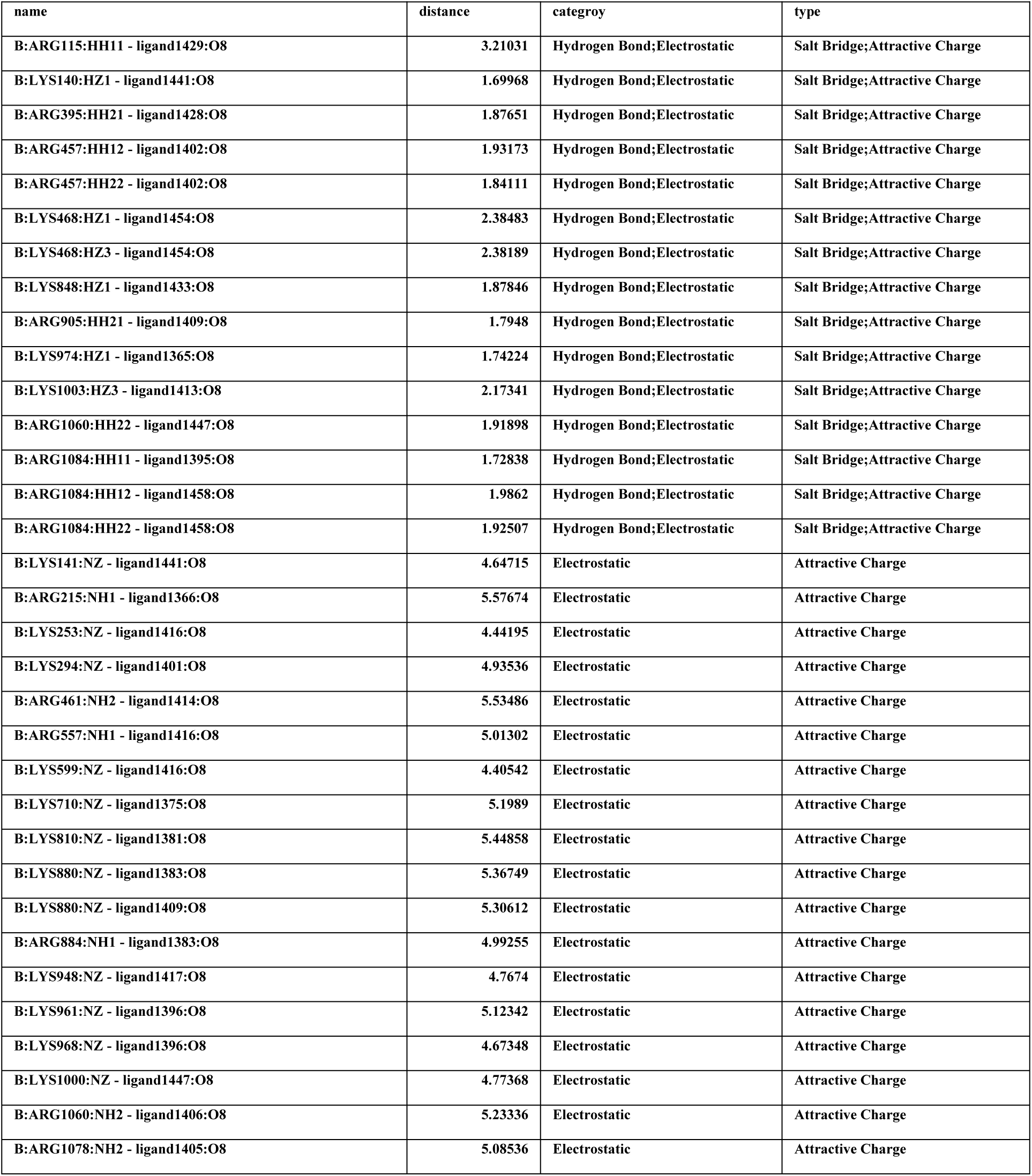
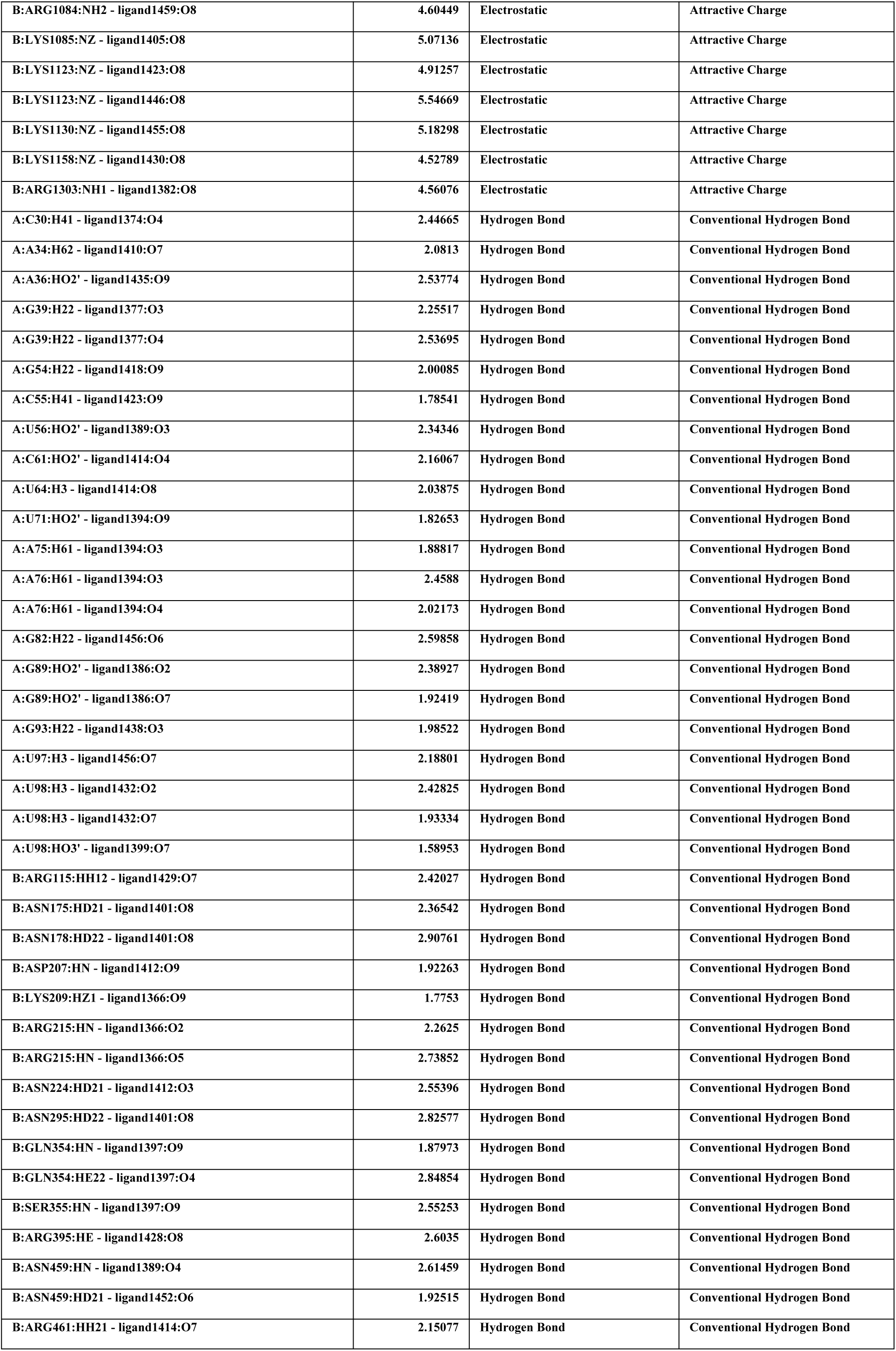
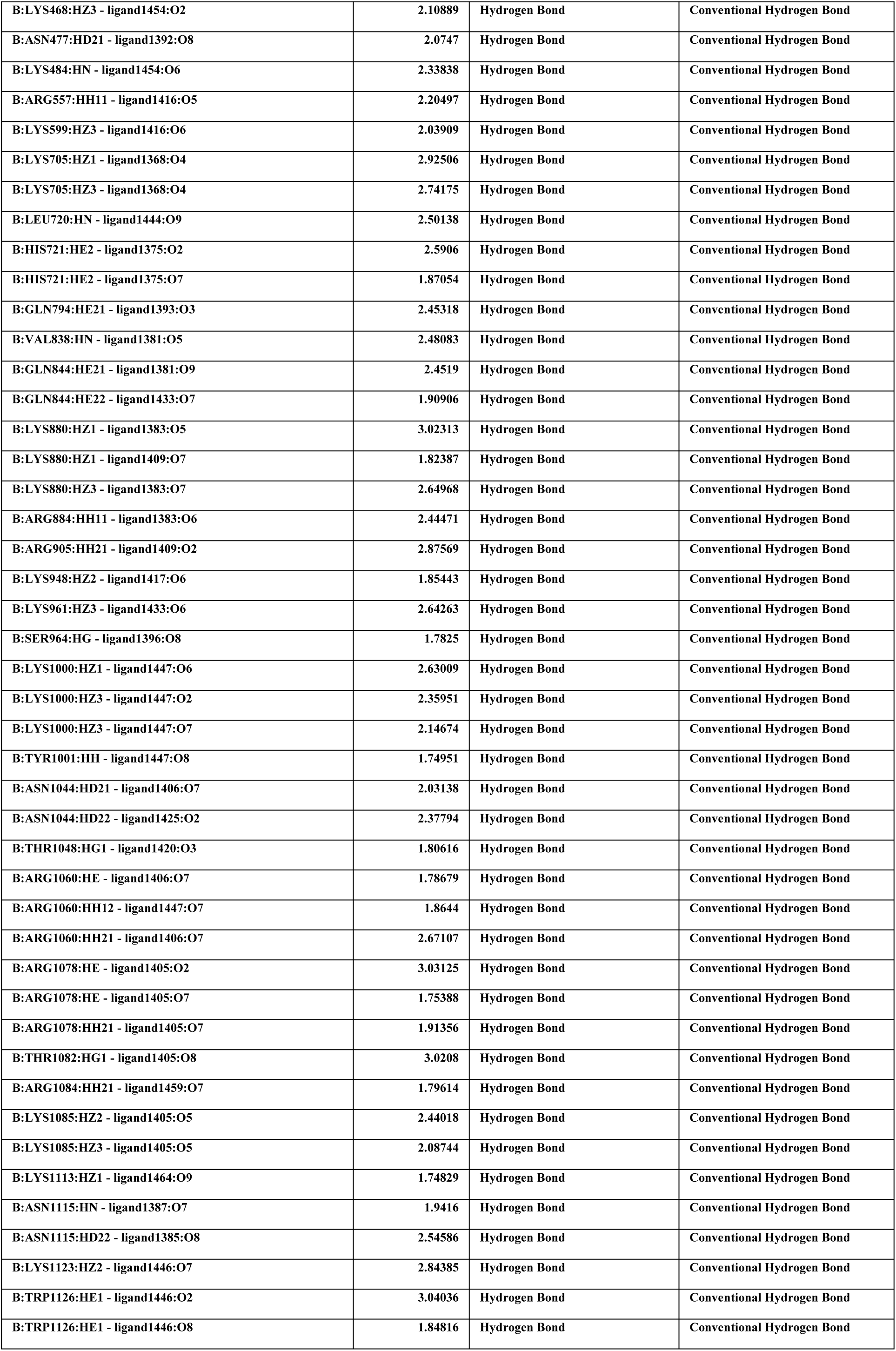
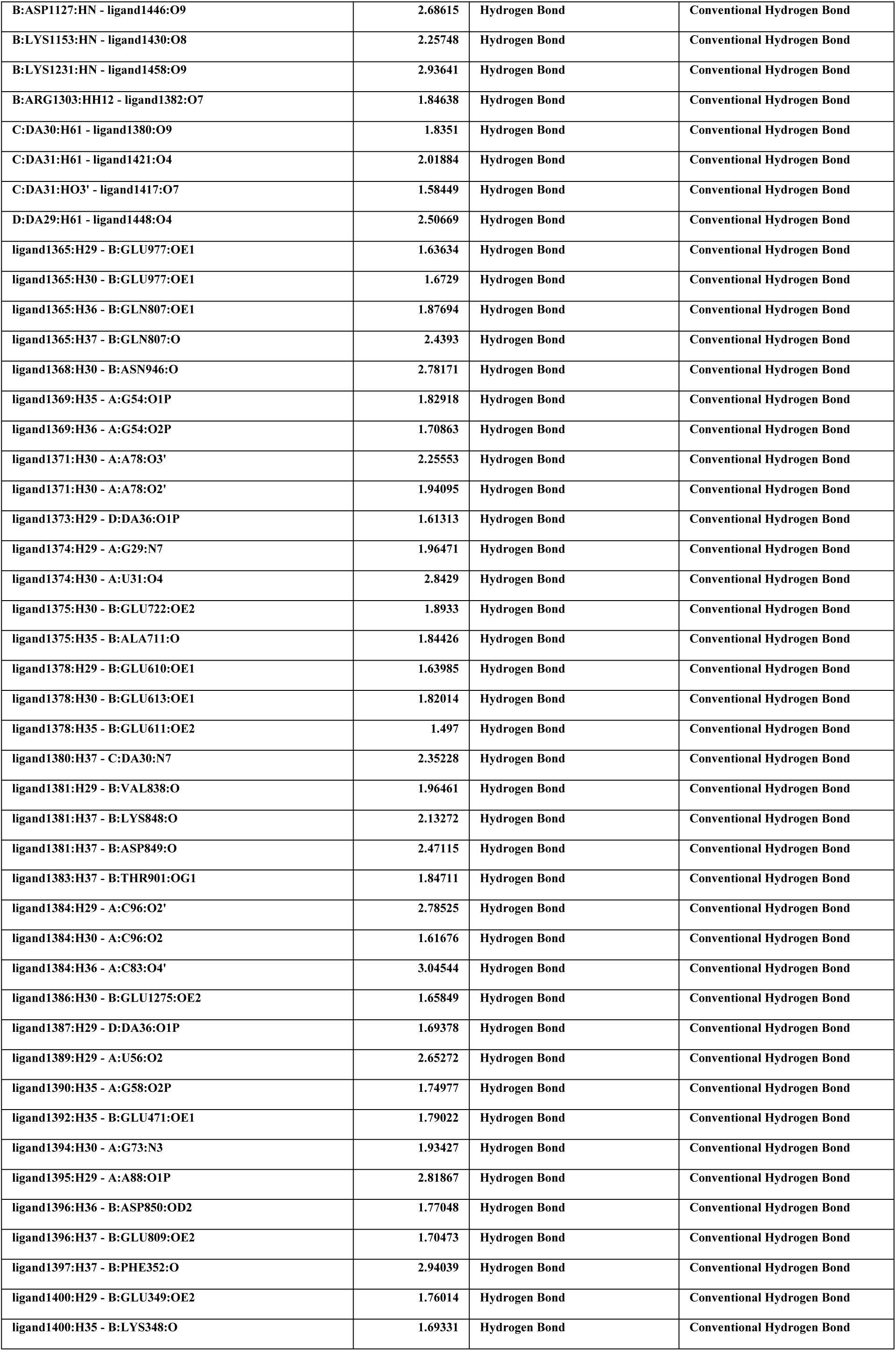
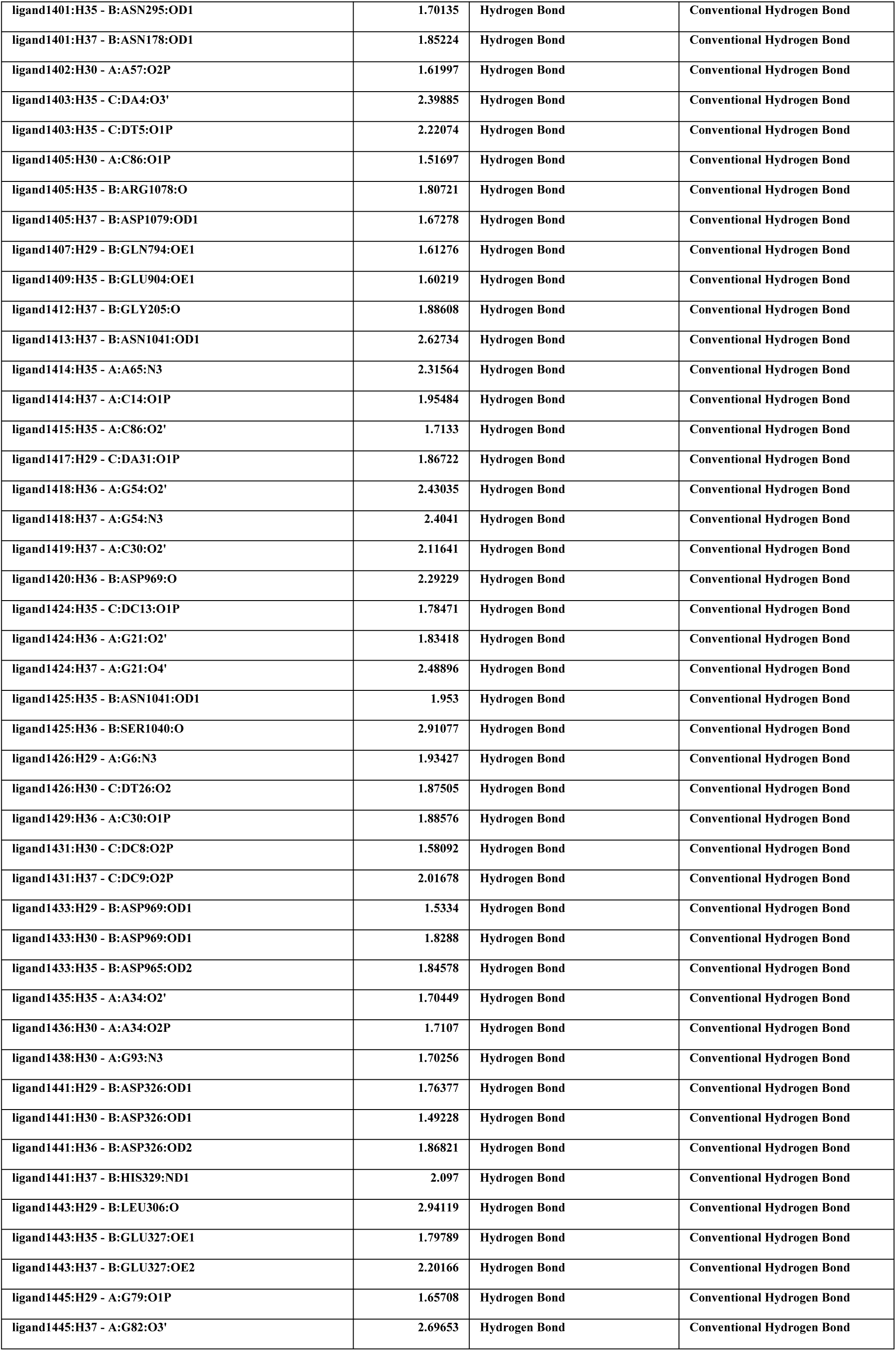
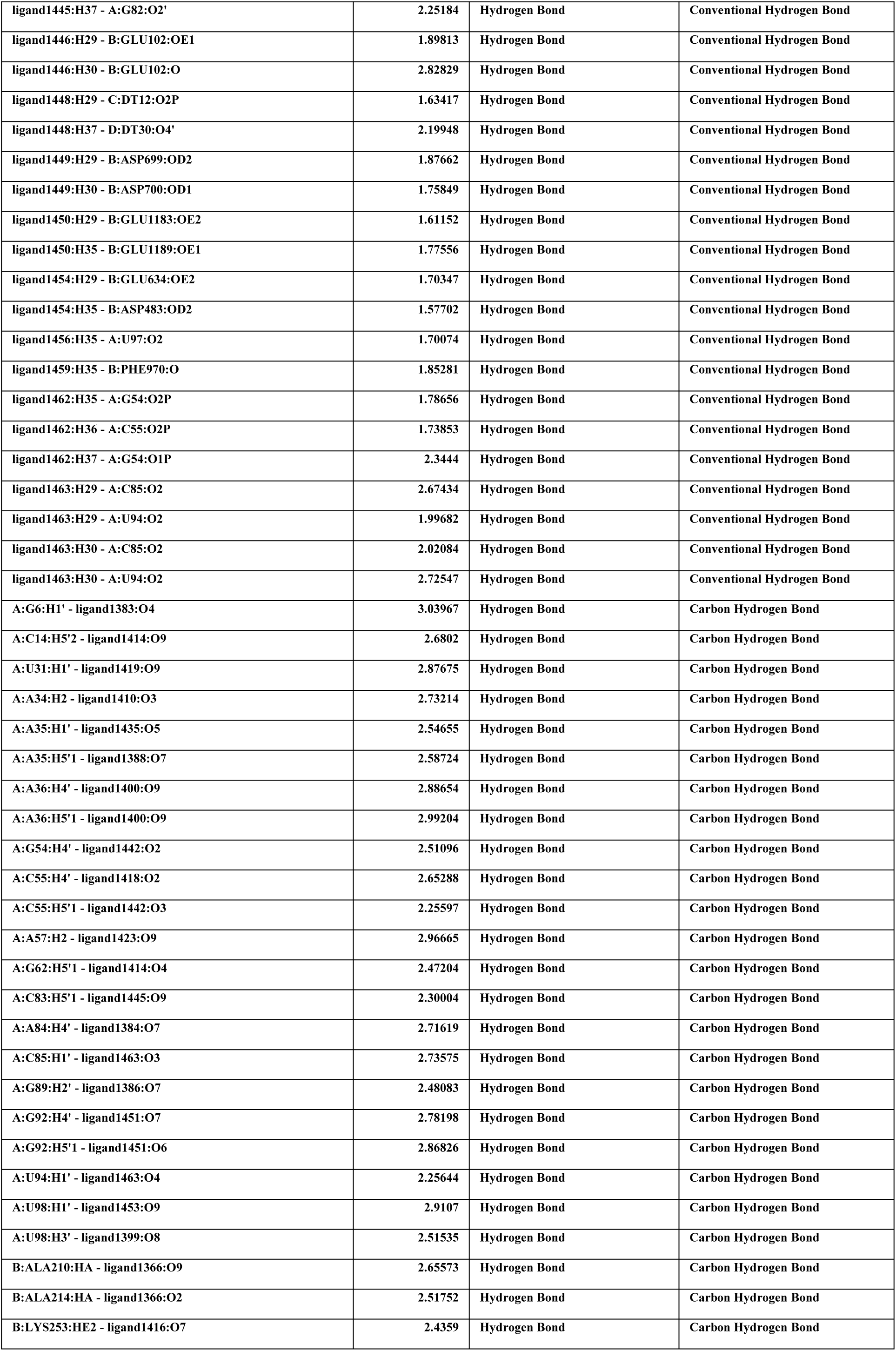
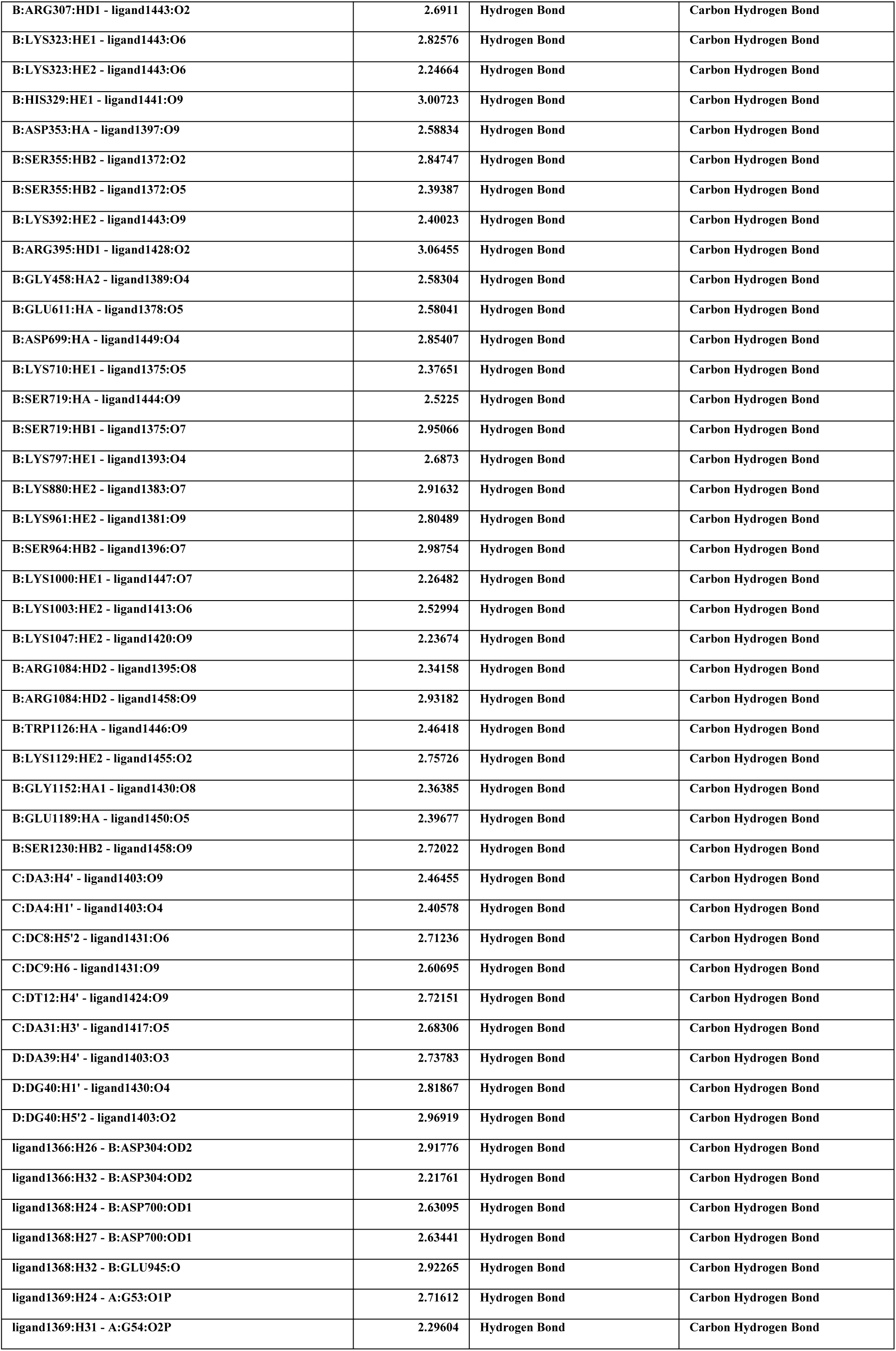
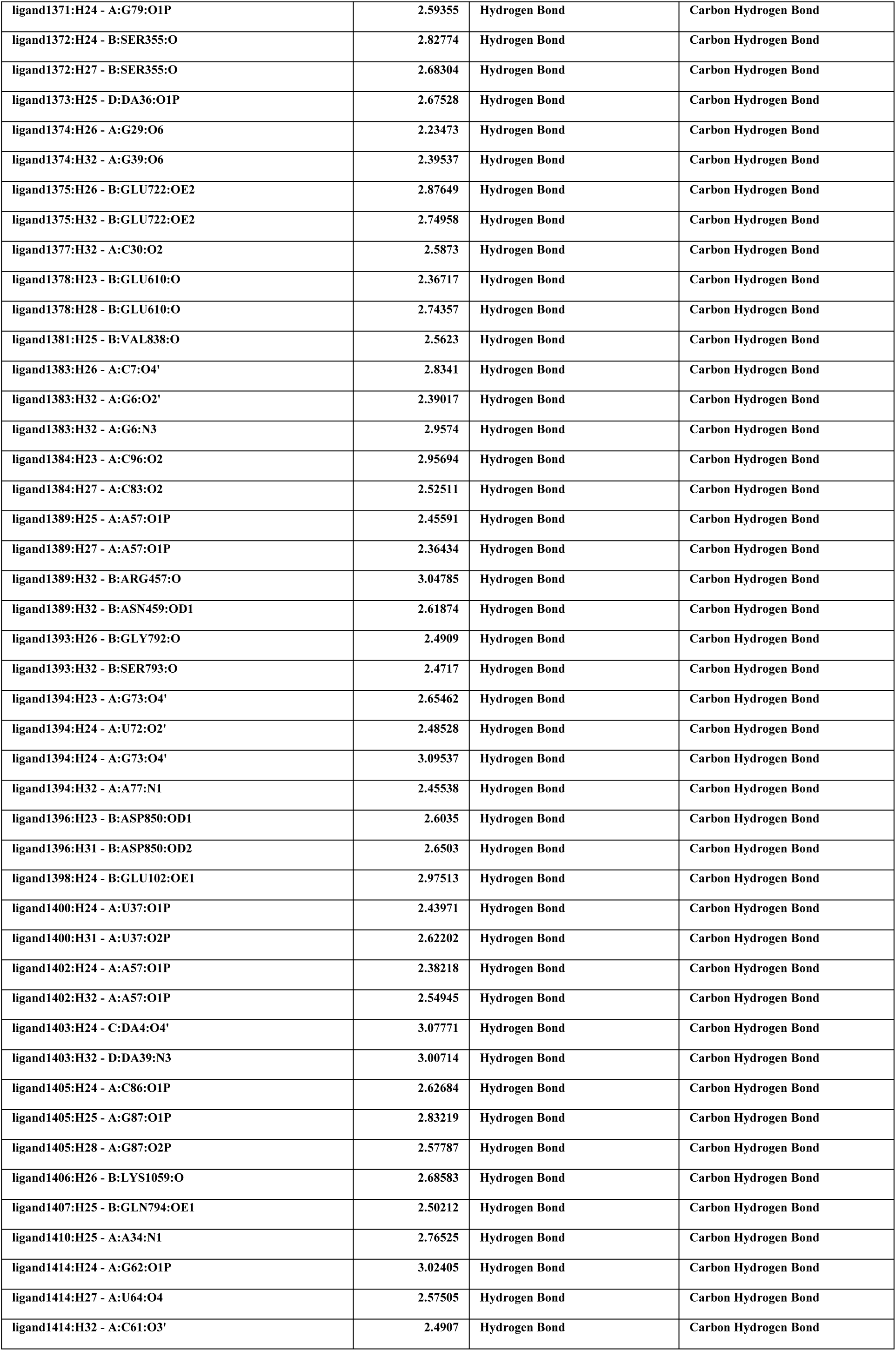
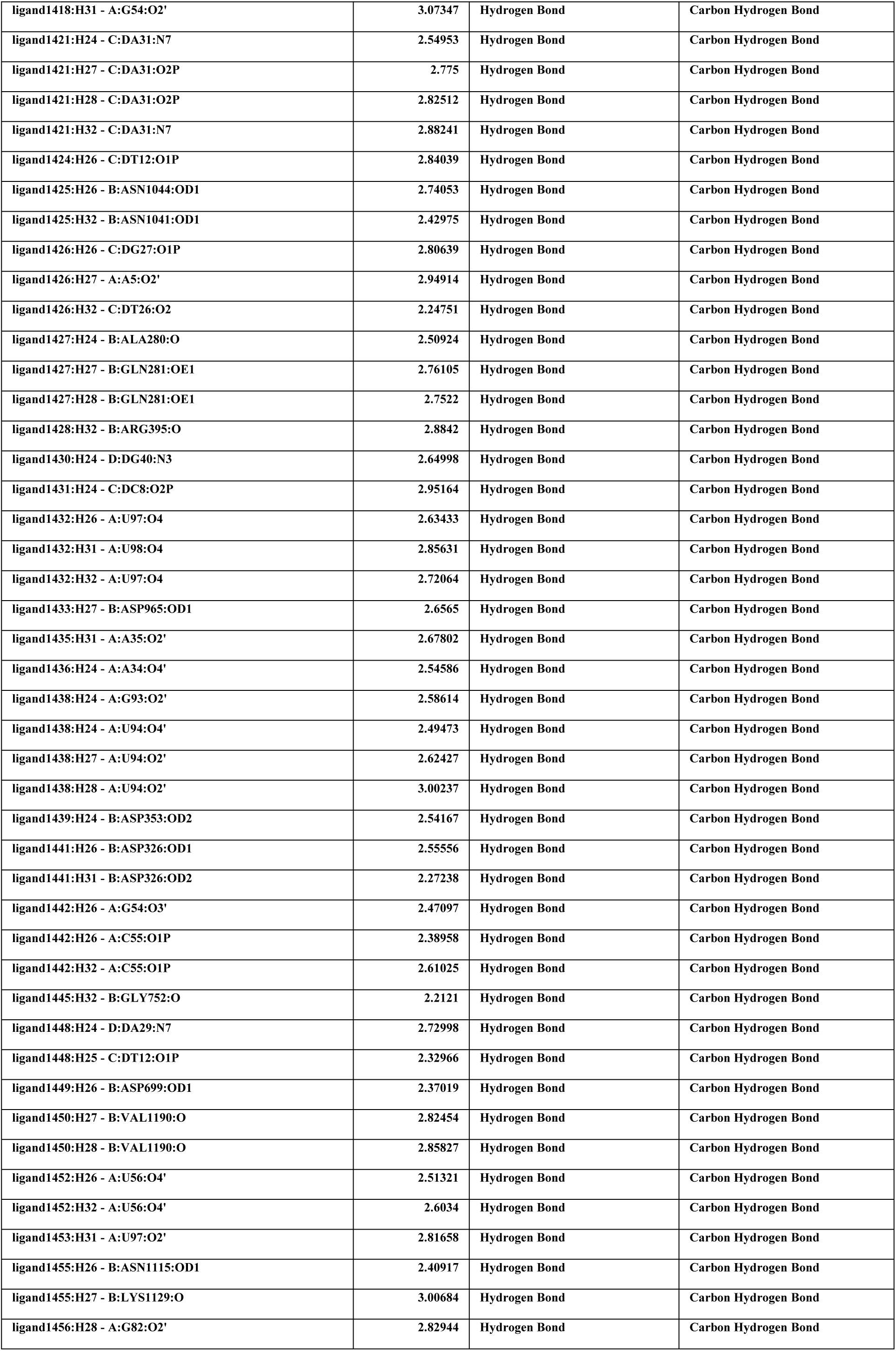
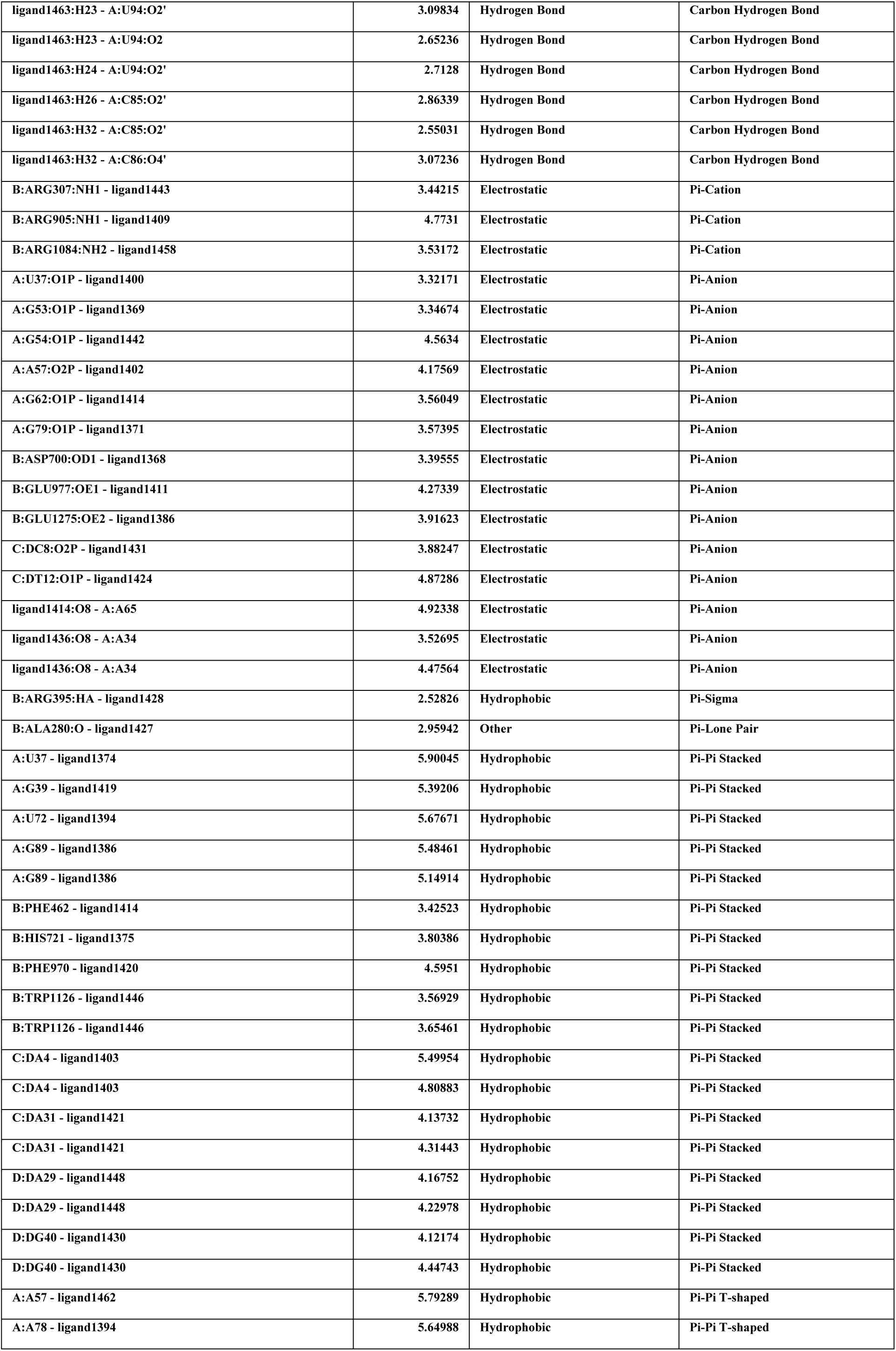
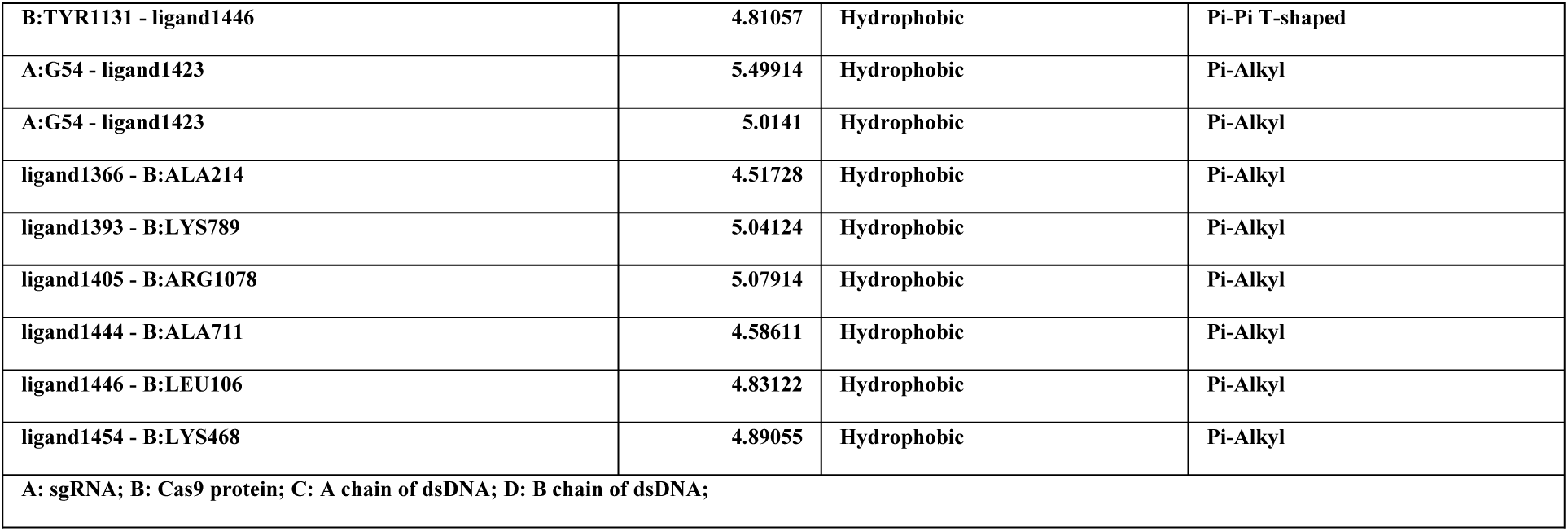
Bonding information of Molecular dynamics of NMN between Cas9 protein and nucleic acids.

**Expand table 3.**
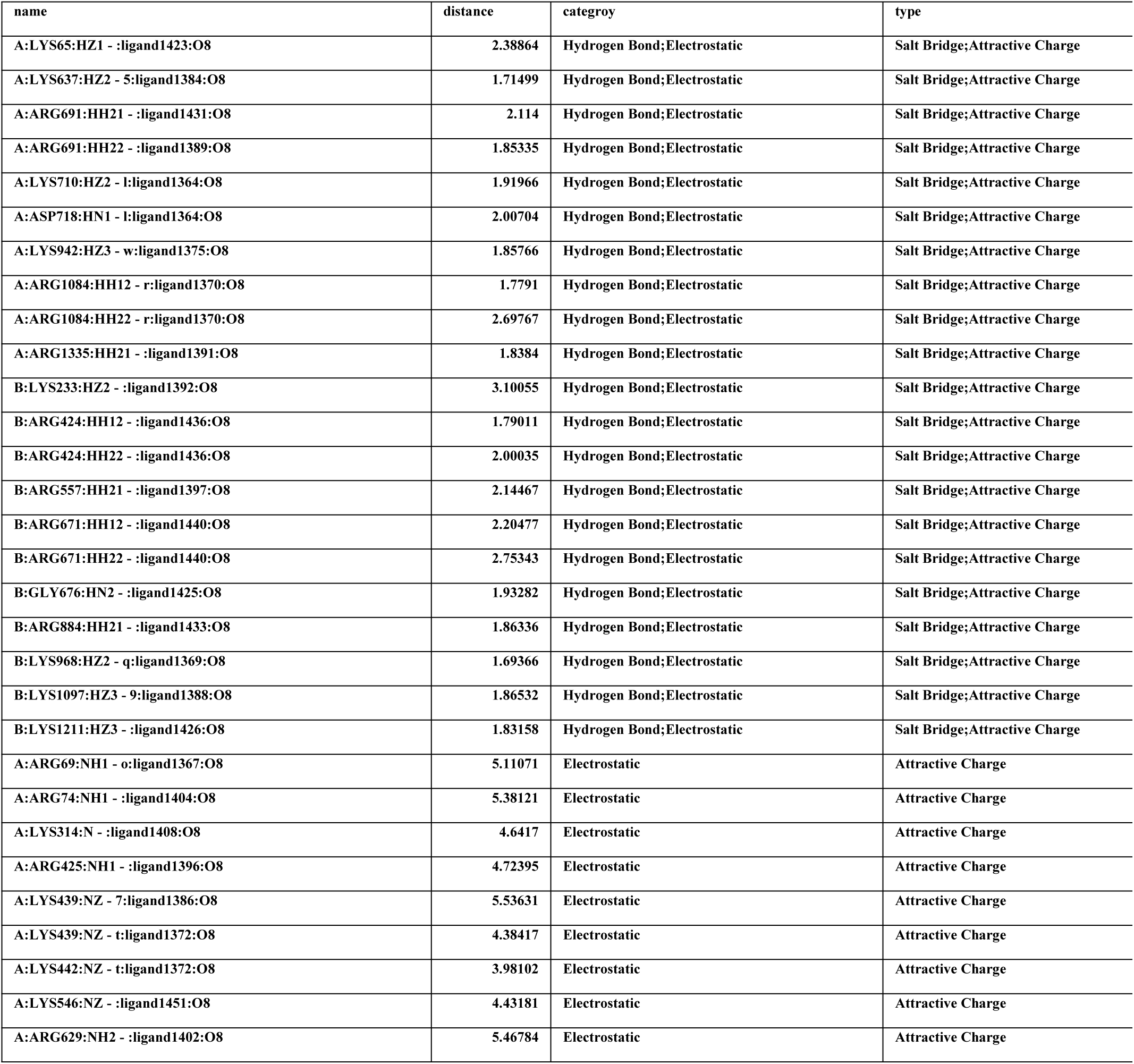
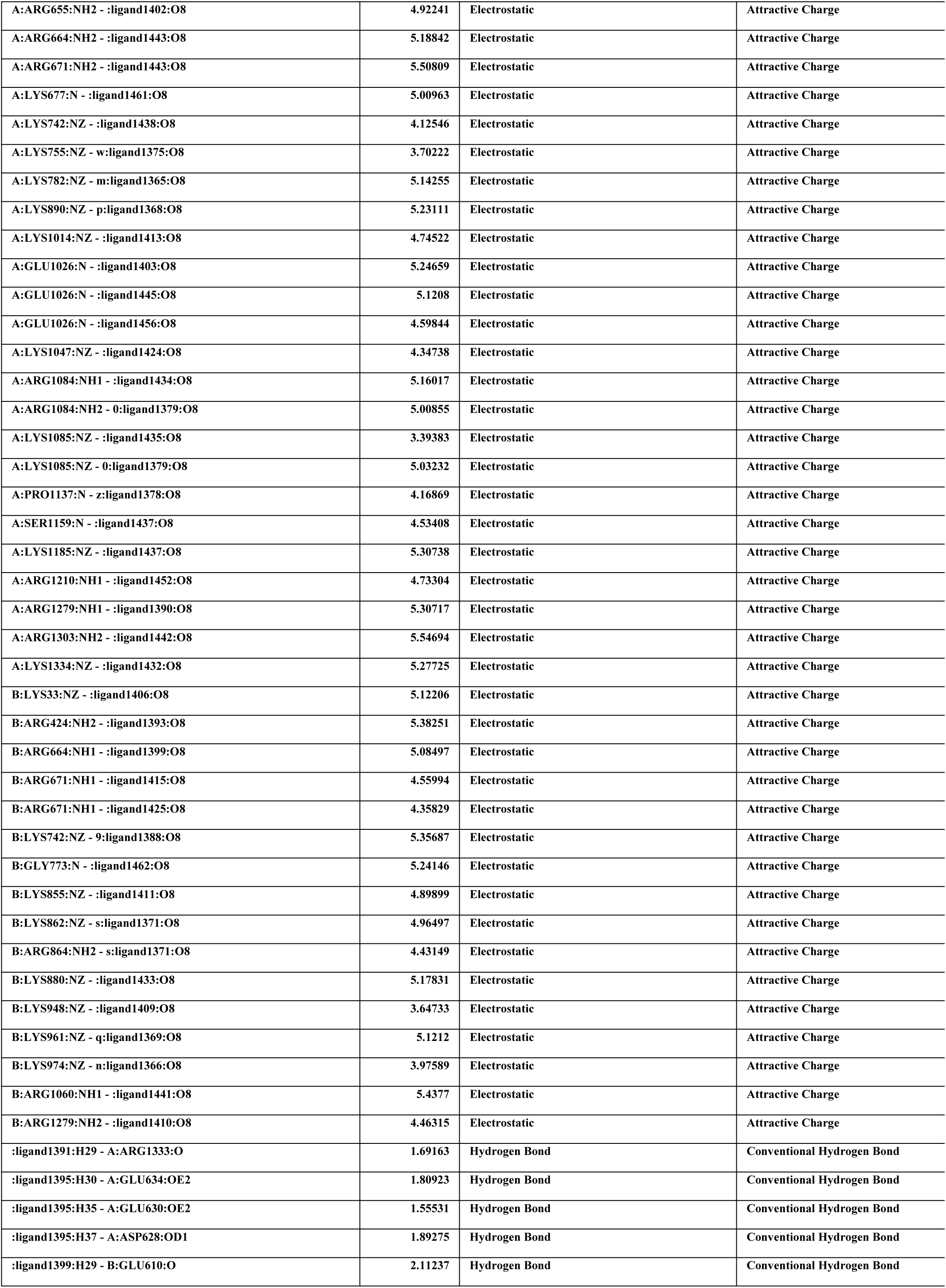
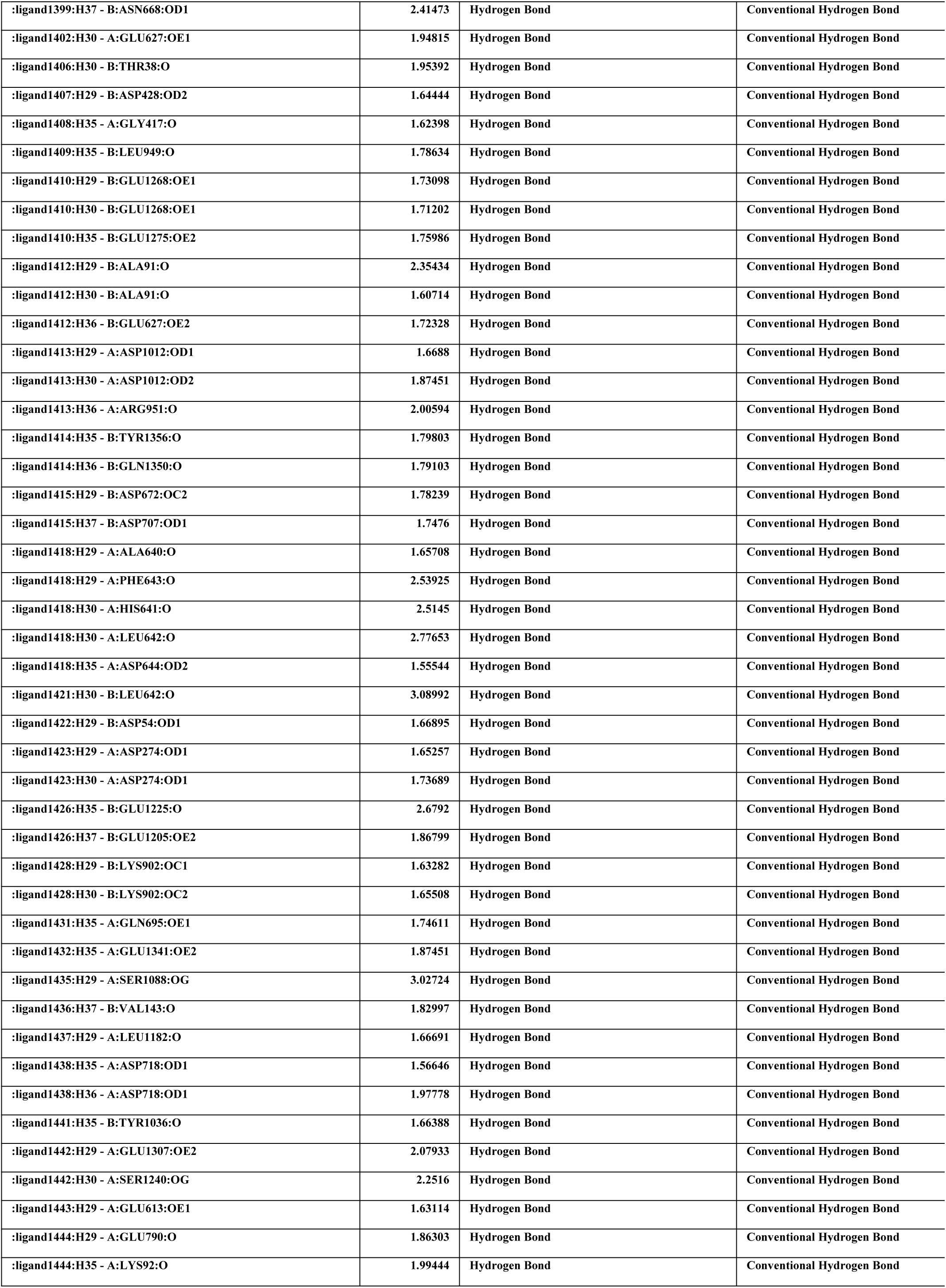
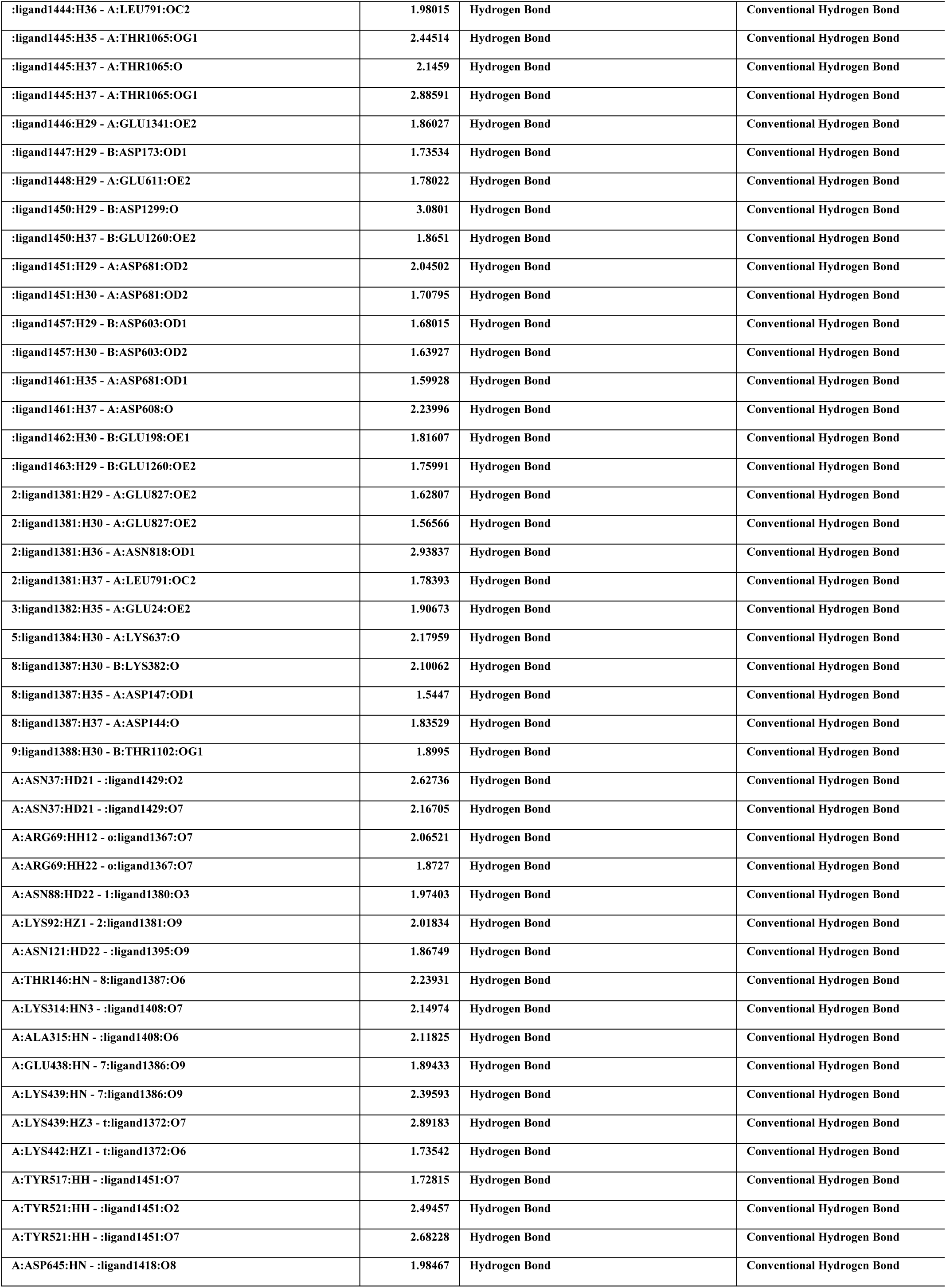
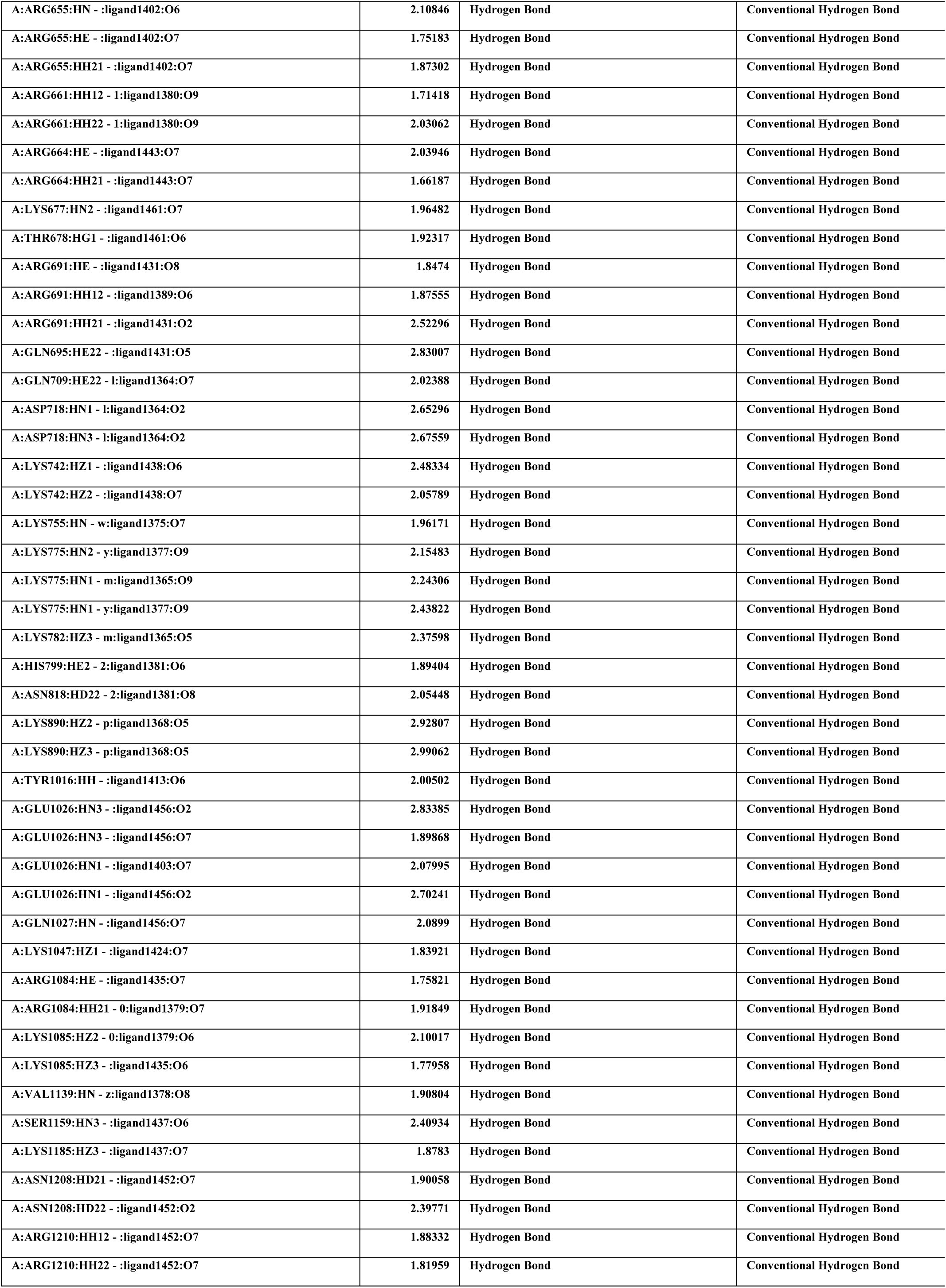
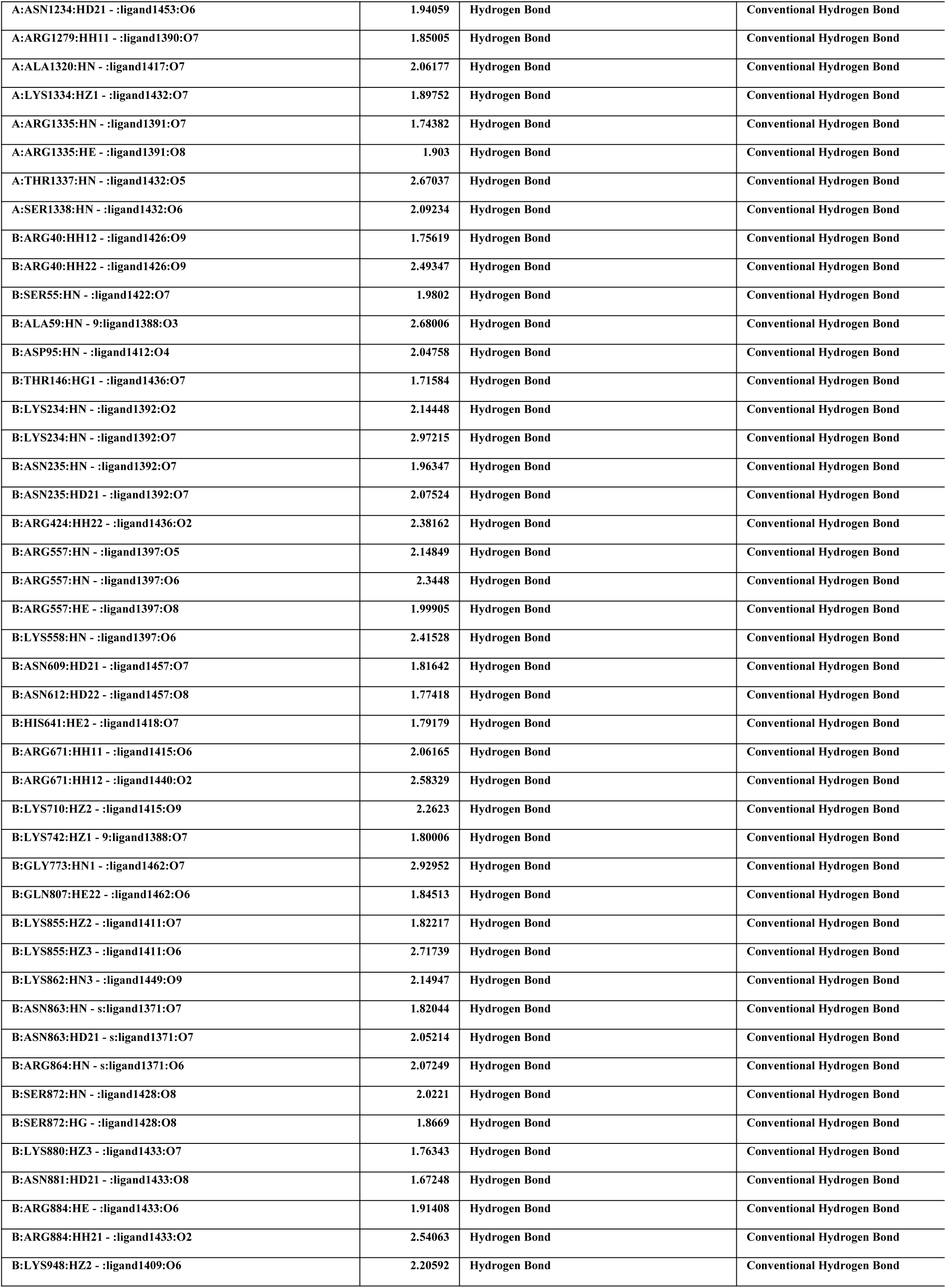
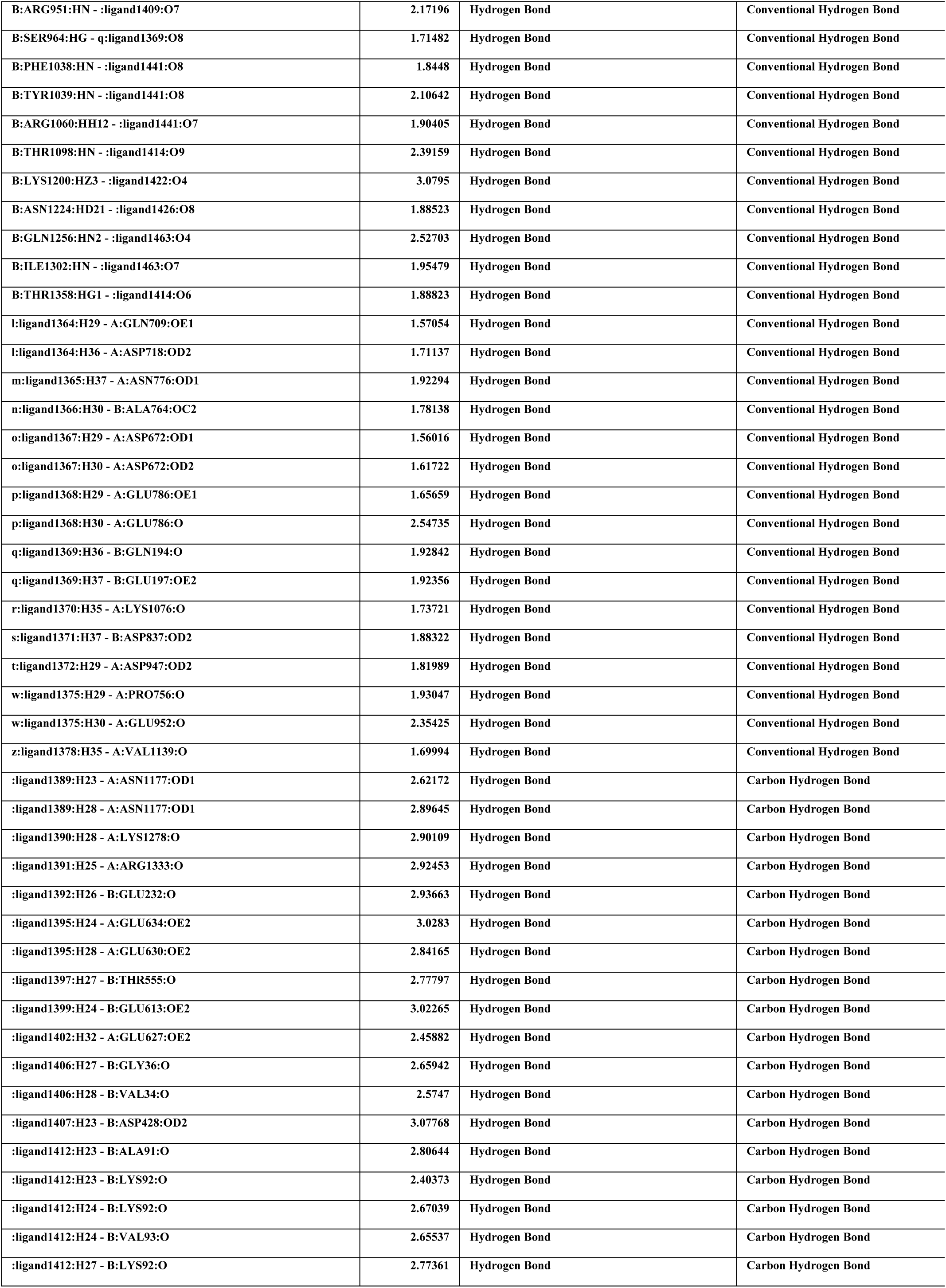
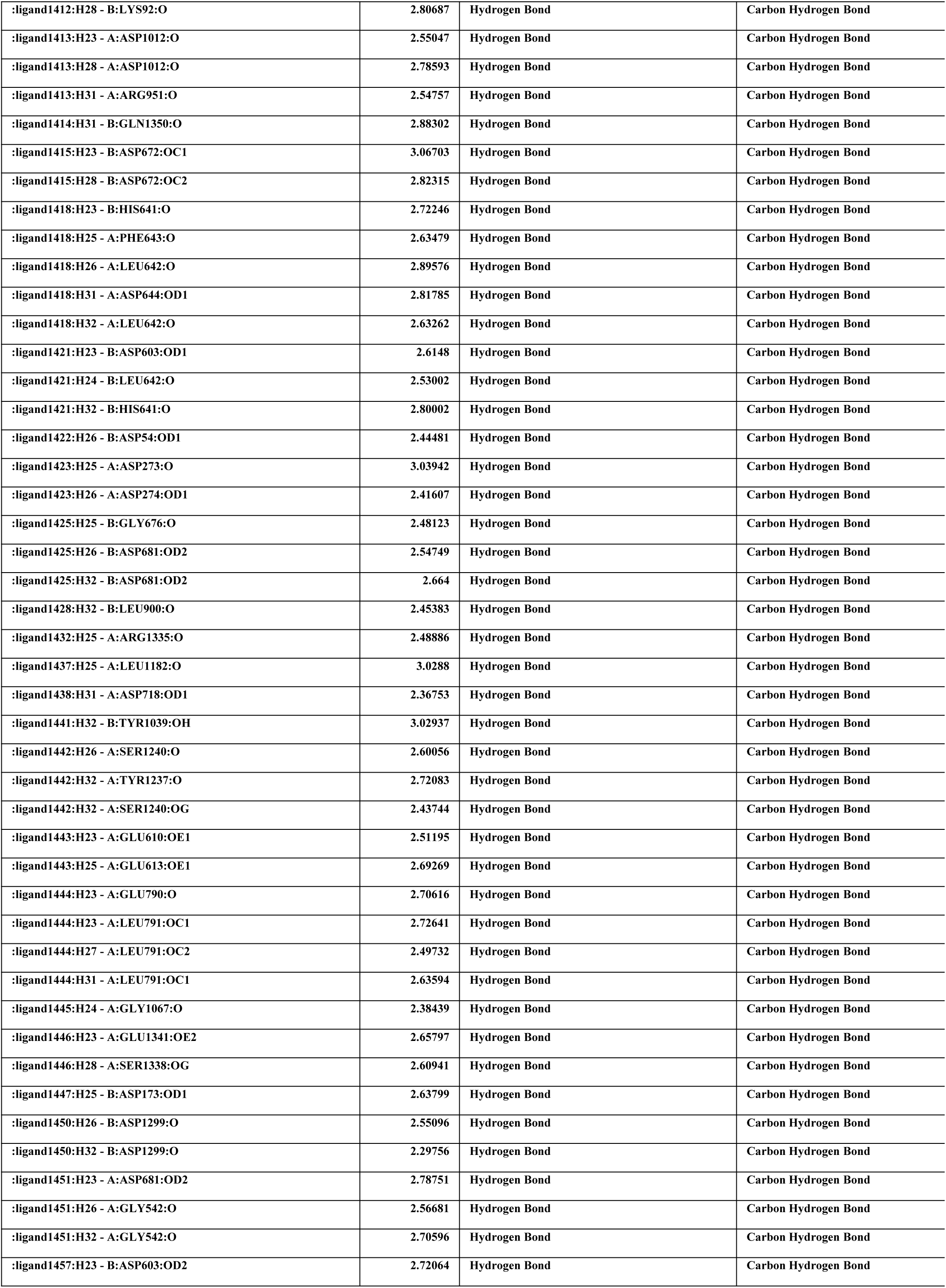
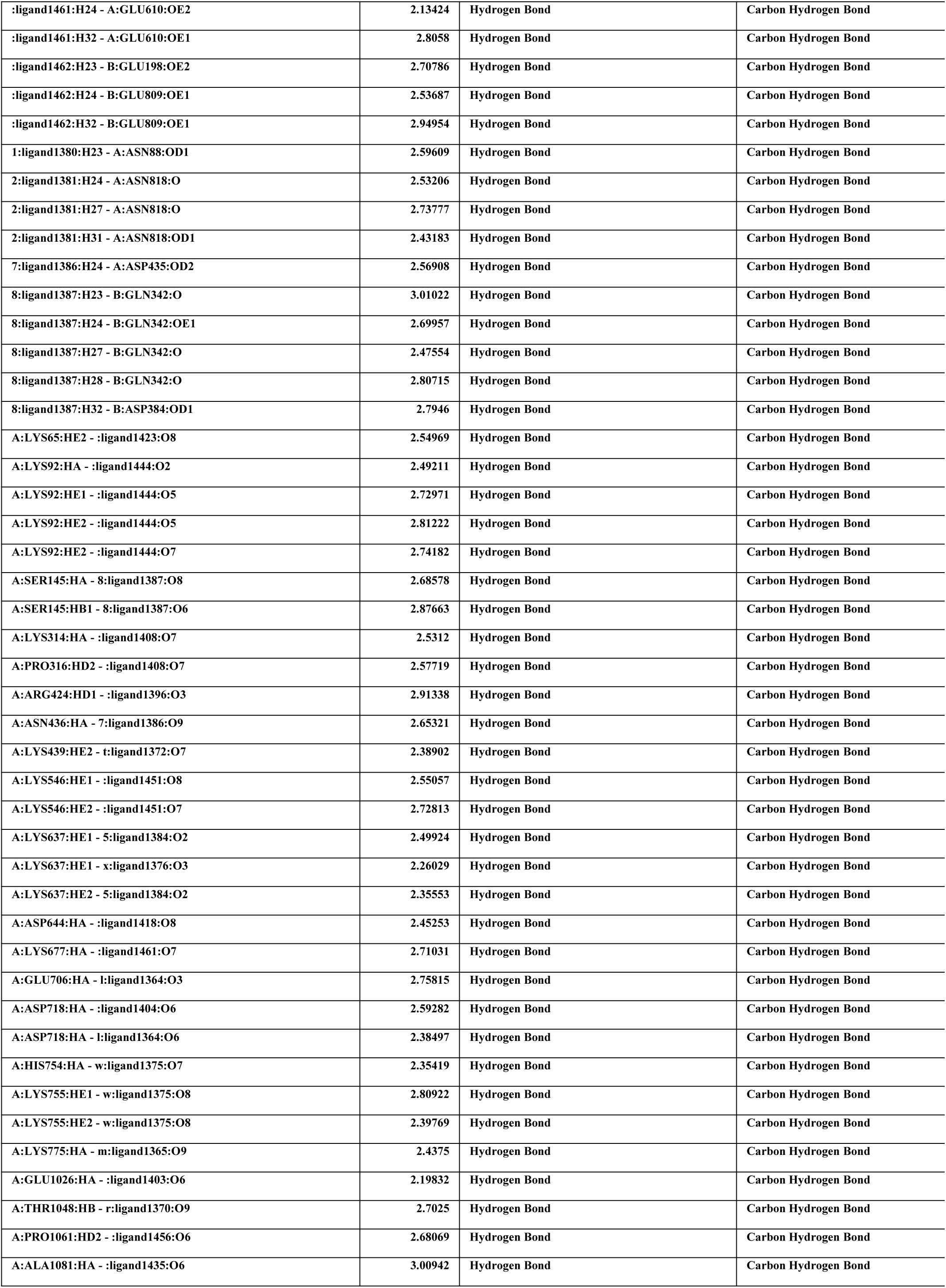
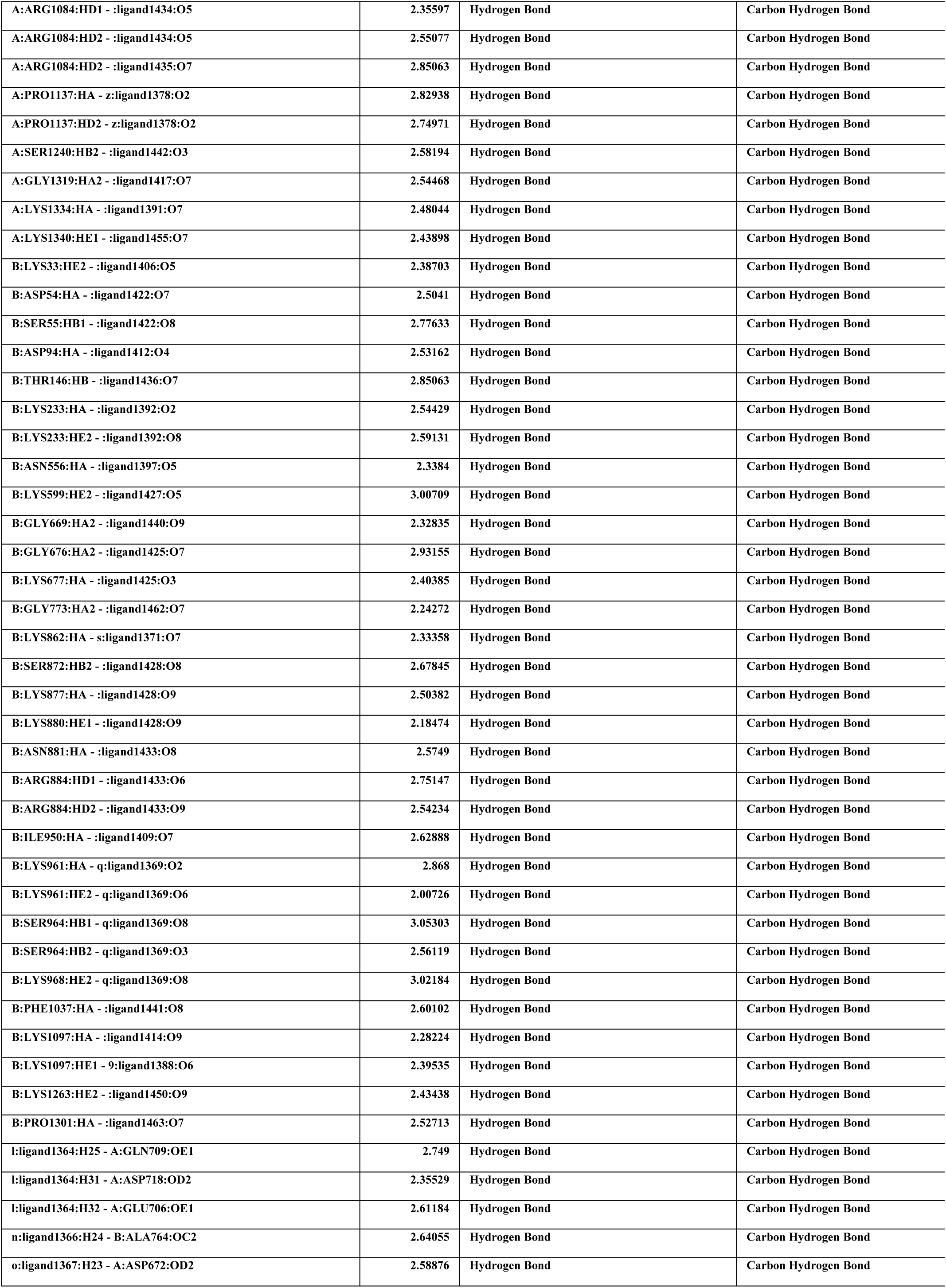
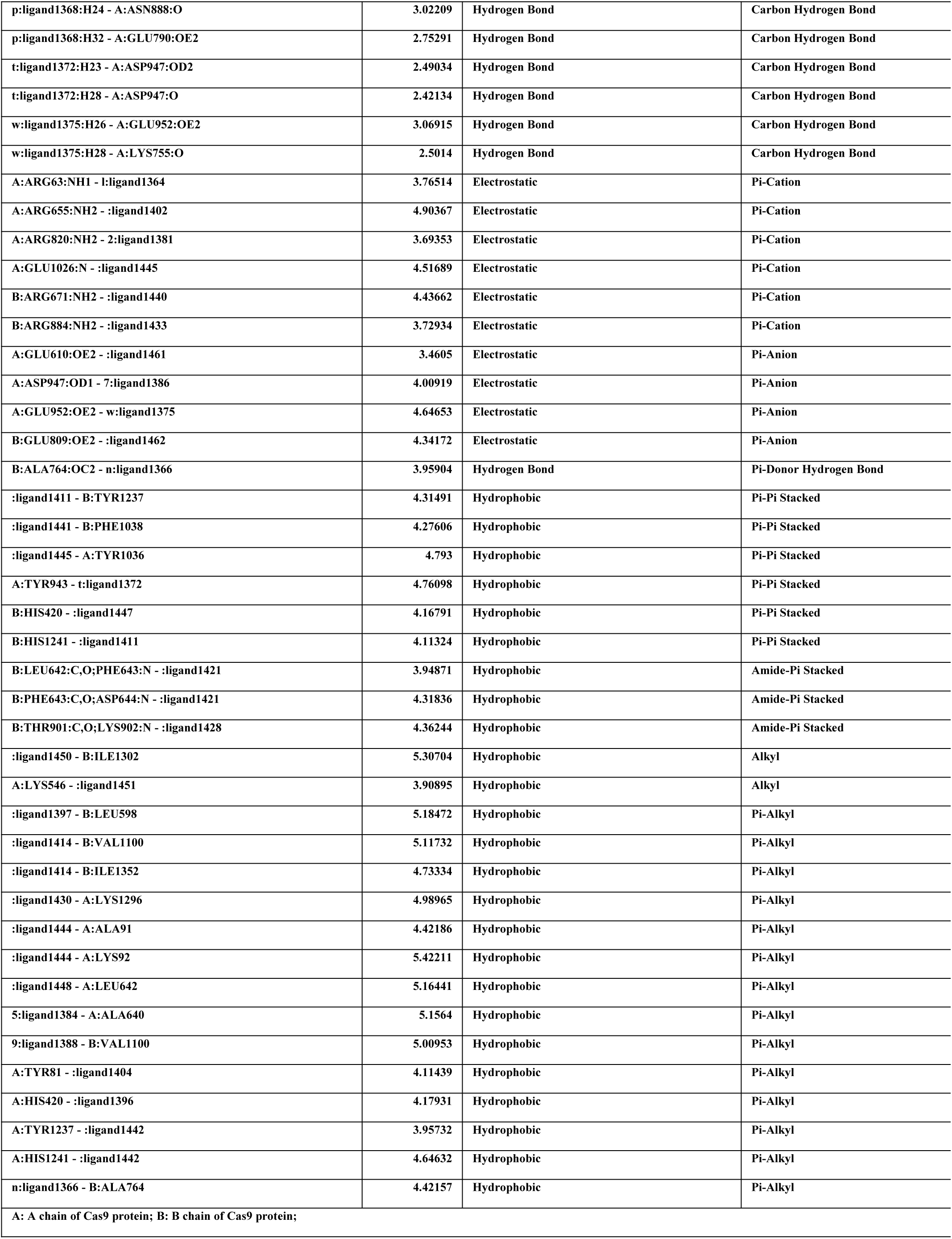
Bonding information of Molecular dynamics of NMN and Cas9 protein.

**Expand table 4.**
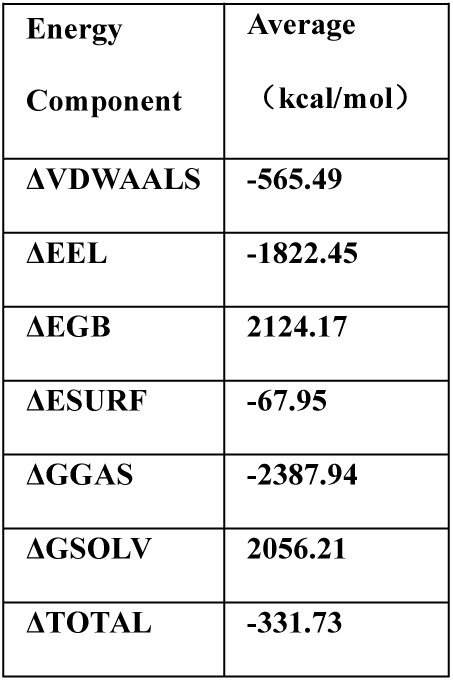
The statistic of combined free energy of NMN and double-stranded DNA.

DNA
| Energy<br>Component | Average<br>(kcal/mol) |
| --- | --- |
| ΔVDWAALS | -565.49 |
| ΔEEL | -1822.45 |
| ΔEGB | 2124.17 |
| ΔESURF | -67.95 |
| ΔGGAS | -2387.94 |
| ΔGSOLV | 2056.21 |
| ΔTOTAL | -331.73 |

**Expand table 5.** The statistic of combined free energy of NMN between Cas9 protein and nucleic acids.

| Energy<br>Component | Average (kcal/mol) |  |  |
| --- | --- | --- | --- |
|  | Protein | RNA | DNA |
| $\Delta$ VDWAALS | -717.04 | -447.59 | -139.43 |
| $\Delta$ EEL | -1546.12 | -1670.81 | -817.41 |
| $\Delta$ EGB | 1964.99 | 1954.34 | 923.44 |
| $\Delta$ ESURF | -111.53 | -50.25 | -16.67 |
| $\Delta$ GGAS | -2263.16 | -2118.41 | -956.84 |
| $\Delta$ GSOLV | 1853.47 | 1904.08 | 906.77 |
| $\Delta$ TOTAL | -409.7 | -214.32 | -50.08 |

**Expand table 6.** The statistic of combined free energy of NMN and Cas9 protein.

| Energy<br>Component | Average<br>(kcal/mol) |
| --- | --- |
| ΔVDWAALS | -1099.56 |
| ΔEEL | -5045.81 |
| ΔEGB | 5366.78 |
| ΔESURF | -183.88 |
| ΔGGAS | -6145.37 |
| ΔGSOLV | 5182.89 |
| ΔTOTAL | -962.48 |

